# On the biological constraints that limit the productivity of rain-fed annual crops

**DOI:** 10.1101/2022.06.21.496884

**Authors:** Iddo Kan, Yacov Tsur, Menachem Moshelion

**Affiliations:** Department of Environmental Economics and Management and The Center for Agricultural Economics Research; The Hebrew University of Jerusalem; Institute of Plant Sciences and Genetics in Agriculture; The Hebrew University of Jerusalem

**Keywords:** transpiration, natural selection, dynamic optimization, wild annual plants, breeding, drought-response optimization

## Abstract

Efforts to cope with hunger by breeding highly productive annual crops for rain-fed agriculture in stochastic-rainfall environments have had only minor success, which we attribute to biological constraints that limit the crops’ yields. We use optimization modelling to interpret experimentally measured transpiration trajectories of wild barley plants following a rain event: the plants first maximized biomass accumulation by employing their maximal transpiration rate, then switched to their minimal transpiration rate to ensure survival until maturity. Thus, breeding plants with lower minimal transpiration rates combined with higher water-use efficiency and maximal transpiration rates could increase expected yields. However, our experimental results indicate that biological constraints impose tradeoffs among maximal and minimal transpiration rates and water-use efficiency. A proposed breeding methodology identifies less biologically constrained cultivar candidates.

## Main Text

Rain-fed annual crops cover nearly 80% of global agricultural land, account for 58% of global agricultural output and are the main source of food for billions of people worldwide (Wani et al. 2009). The increased demand for food (Tilman et al. 2011) and crop sensitivity to climate change (Lobel et al. 2014, Zhao et al. 2017), together with projections for a low rate of expansion of irrigated lands (FAO 2017), underscore the need to increase yields of rain-fed crops, particularly in regions in which rainfall timings and levels fluctuate widely. Conventional breeding programs aimed at identifying high-yield phenotypes via field experiments are slow and costly because of the interactions between dynamic soil-atmosphere conditions and the enormous number of genes and germplasm candidates that can potentially improve yield (Negin and Moshelion 2017).

Recently developed methods for accelerating breeding employ advanced biotechnologies (Takeda and Matsuoka 2008, Skirycz et al. 2011, Millet et al. 2019) and pre-field screening measures (e.g., delayed leaf senescence, photosynthesis, water-deficit recovery rates and stomatal conductance). Nevertheless, as there is a wide genotype-to-phenotype knowledge gap (Tuberosa et al. 2014) and genome×environment×management interactions are not fully understood (Großkinsky et al. 2015), there is no consensus in the literature as to how to proceed from data to knowledge (Blum 2009, Moshelion and Altman 2015) and the bench-to-field transfer rate of commercial drought-tolerant crops remains low (Graff et al. 2013, Dalal et al. 2017). We propose to mitigate this shortcoming by using evolutionarily selected (revealed) traits of annual wild plants as benchmarks in the breeding of highly productive rain-fed crops under dry conditions.

Rain-fed cultivars have inherited the traits of their wild ancestors (Mickelbart et al. 2015), which evolved through natural-selection processes that favored the most fit (Volis et al. 2004). As fitness and reproduction (i.e., grain generation) are positively correlated (Sultan 2000), traits that maximize the fitness of wild plants for specific environments should closely match those that maximize the expected yields of their domesticated rain-fed descendants under similar conditions. Therefore, understanding the principles underlying the natural selection of physiological traits of wild plants can provide useful guidance for breeders of rain-fed crops. We focus on the evolution of plant traits that dictate the patterns of the transpiration behavior (hereafter *transpiration policy*) employed by wild annual plants in environments with uncertain rainfall.

Optimizing water management is a crucial aspect of the survival of wild terrestrial plants. Since the pioneering work of Cowan and Farquhar (1977), many scholars have adopted the concept that plants maximize carbon gains subject to water-availability constraints (e.g., Mencuccini et al. 2019). To the best of our knowledge, only two studies have explicitly incorporated uncertain rainfall in their optimization models: Mäkelä et al. (1996), who characterized optimal transpiration trajectories following a single rainfall event, and Lu et al. (2016), who considered an infinite number of consecutive rainfall and dry-down events. However, both of those approaches are inappropriate for analyzing transpiration by annual plants, because they overlook the plant’s maturity (flowering) time, which is the quantitative trait most adaptive to dry conditions (Verhoeven et al. 2008, Richards et al. 2010). Here, we characterize the link between the transpiration policy and maturity time by integrating insights from a theoretical model with results from a large experimental data set of continuous and simultaneous soil-plant-atmosphere continuum measurements of multiple wild-barley accessions.

We view the natural selection of wild annual plants as a two-stage optimization process (Fig. 1). In the first stage, evolutionary processes have optimized the plants’ transpiration policy subject to two sets of factors. The first set of factors includes the environmental conditions in the plant’s habitat, such as the frequency and intensity of rain events, vapor-pressure deficit (VPD), soil structure and neighboring vegetation. The second set incorporates four plant physiological traits: maturity time, water-use efficiency and the maximal and minimal transpiration rates. Experimental observations indicate that, during dry-down events, the transpiration trajectories of wild barley plants switch from high to low rates of transpiration (Galkin et al. 2018). Our formal analysis interprets this transpiration behavior as the plant’s optimal transpiration policy under stochastic rainfall conditions. This policy maximizes the plant’s expected biomass at maturity, as the plant switches from maximal to minimal transpiration at the precise time that avoids drying out exactly until it reaches maturity.

**Fig. 1.**
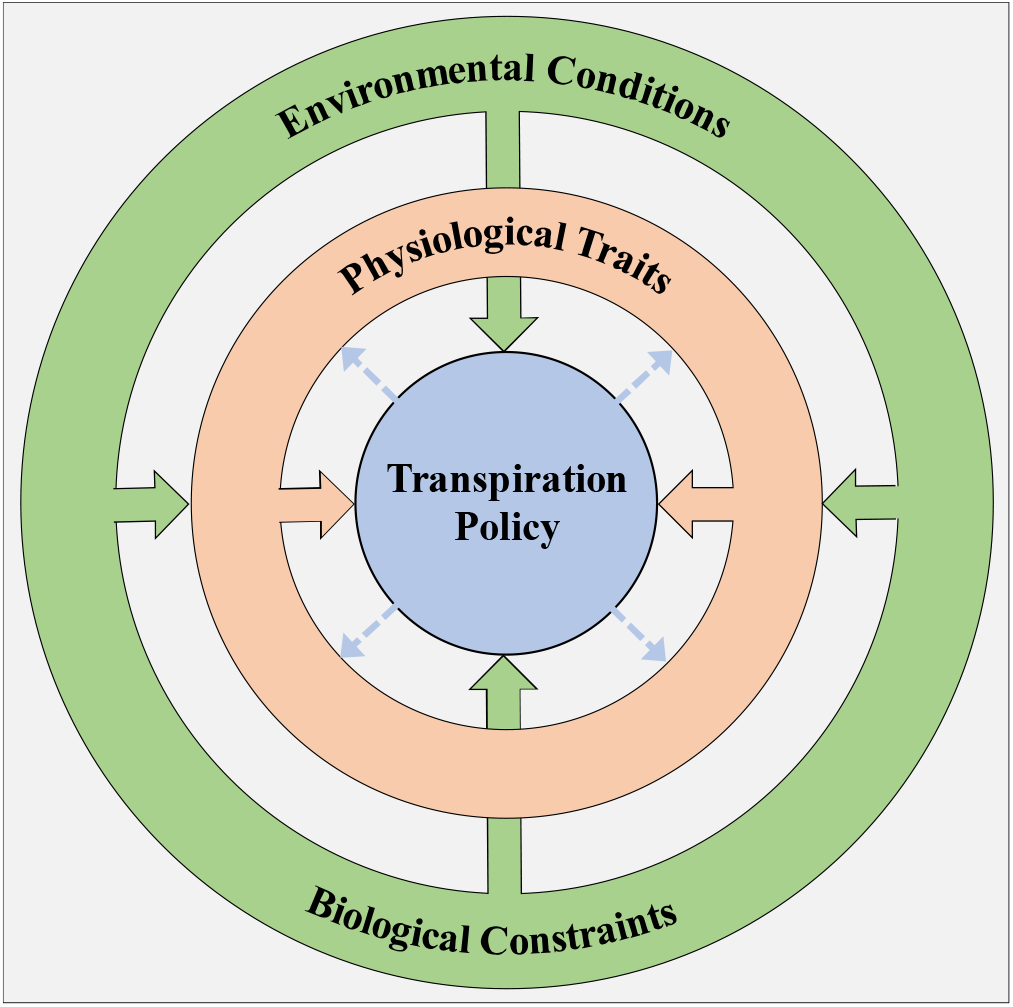
Schematic hierarchy of transpiration and the selection of physiological traits. Environmental conditions and biological constraints (outer ring) affect the selected physiological traits of a specific genotype (middle ring). Those traits, along with the environmental conditions, dictate the genotype’s transpiration policy (inner circle). The dashed-line arrows indicate that the selection of the physiological traits occurs in accordance with the patterns of the transpiration policy.

In the second optimization stage, natural-selection processes have selected the set of physiological traits that maximizes the efficacy of the optimal transpiration policy of wild plants in relation to the specific environmental conditions in their habitats. We show that, as a general rule, natural-selection processes should have favored plants with greater water-use efficiency and maximal transpiration rates (and, therefore, higher rates of biomass accumulation) combined with lower minimal transpiration rates—the latter enable survival until prolonged maturity times and thereby the utilization of more rainfall events for biomass accumulation throughout the growing season. However, our empirical measurements of the traits of wild barley plants indicate that water-use efficiency and the maximal and minimal transpiration rates are biologically linked with one another in a way that imposes tradeoffs between these traits. We, therefore, conclude that maturity time is the most adaptive trait for dry conditions because it is relatively biologically independent of the other traits that dictate the optimal transpiration policy. Our proposed methodology for breeding annual rain-fed crops for production in a specific environment favors germplasm candidates whose traits are less biologically constrained relative to the naturally (i.e., optimally) selected traits of the candidates’ wild ancestors that evolved in similar environments.

This paper is structured as follows. In the following three sections, we deal with the inner circle of Fig. 1 and characterize the optimal transpiration policy. Then, we model the natural selection of the four physiological traits (middle ring of Fig. 1) as a constrained optimization problem. After that we present our proposed breeding methodology and, in the final section, we present our conclusions. Appendices A and B provide technical detail regarding the theoretical analyses (Appendix A) and the empirical analyses (Appendix B).

### Observed Transpiration Patterns

We used data from the B1K collection (Hübner et al. 2009, Hübner et al. 2013) of wild-type barley (*Hordeum vulgare* sps. *spontaneum*) for barley specimens that are native to five different sites across Israel (Meron, Arbel, Oren, Guvrin and Yeruham). Those sites all have distinct environmental conditions and the examined lines exhibited a high degree of genetic variation (Galkin et al. 2018). In Fig. 2, we present results from a water-deficit pot experiment applied to a plant from Arbel (all accessions exhibited qualitatively similar responses), using the PlantArray high-throughput functional-phenotyping gravimetric system (Galkin et al. 2018). In Fig. 2A, we present time trajectories of daily average transpiration per unit of leaf area (*E*, measured in mmol sec^-1^ m^-2^) and VPD (kPa) as the atmospheric force driving transpiration, from the time the irrigation was turned off (29 days after planting) onward. The associated trajectories of soil water content (SWC) at the soil volume attainable by the plant’s roots (denoted *θ*, mmol m^-2^ and presented in Fig. 2B in relative terms: 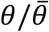, where 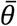 is the maximal SWC, for example, at field capacity) and leaf area (LA, m^2^) is a measure of the plant’s biomass (denoted *m*).

**Fig 2.**
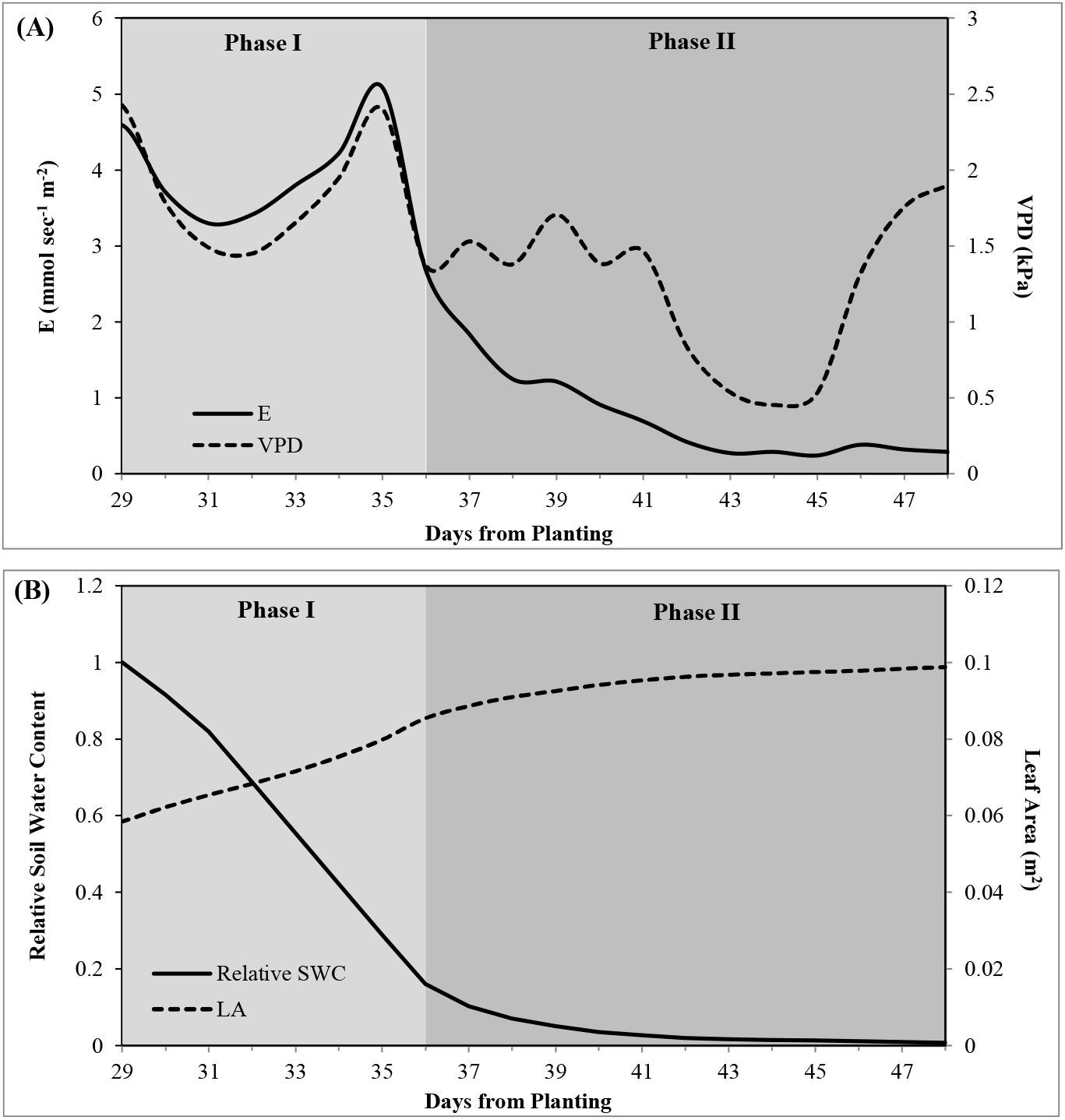
A biphasic transpiration path observed during a water-stress pot experiment applied to a wild barley plant from the Arbel site of the B1K collection (Galkin et al. 2018). Irrigation was stopped at 29 days after planting. (A) Daily average transpiration *E* and VPD. (B) Relative soil water content 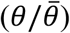 and leaf area as a measure of biomass (*m*).

We can see that transpiration, *E*, evolves along a biphasic trajectory. During Phase I, *E* is high and varies with VPD. During this phase, biomass (LA) increases rapidly and the SWC declines quickly. During Phase II, transpiration proceeds at a low rate and is less dependent on fluctuations in VPD; correspondingly, the rate of biomass growth and the rate at which relative-SWC declines are both lower. This biphasic transpiration behavior is characteristic of annual plants (Moshelion et al. 2015, Negin and Moshelion 2017) and we consider this to be the optimal transpiration policy employed by the plants under uncertain rainfall conditions.

### The Transpiration Dilemma

Consider a seasonal wild plant in an environment with sporadic rainfall events of uncertain intensity (Fig. 3). Germination takes place at the onset of a rain event, initiating a growing season of a fixed (predetermined) length until maturity (i.e., flowering; Merchuk-Ovnat et al. 2018), referred to as 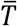. During a growing season, there is a random number of rainfall events, each of which replenishes the SWC in the root zone, marking the beginning of a new management period, which lasts until the next rain event, maturity or desiccation, whichever comes first.

**Fig 3.**
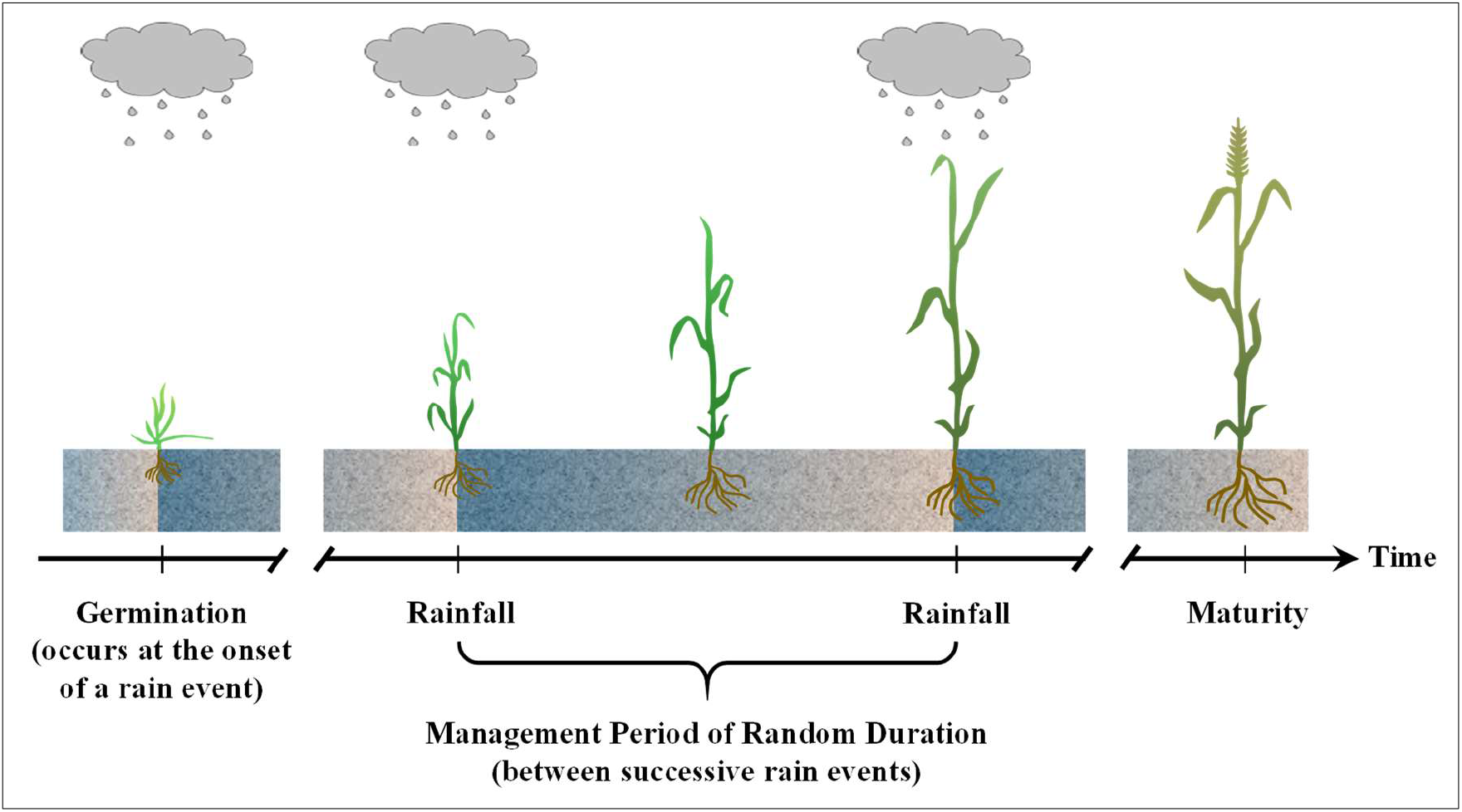
Germination occurs at the onset of a rain event, initiating a growing season of length 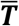. The growing period includes a series of management periods of uncertain duration, each initiated by a rainfall event. A rainfall event replenishes the SWC, which then decreases throughout the management period until the next rainfall event, and so on.

Let *t* denote the time from the onset of some management period *i, i* = 1,2, ….. During the management period, the SWC steadily declines as the soil loses water to the plant’s own transpiration, as well as to direct evaporation and the uptake of water by adjacent plants. We refer to the water loss that is not due to transpiration as drainage (denoted *D* and measured in the same per-biomass units as transpiration; i.e., mmol sec^-1^ m^-2^) and note that *D* depends on the VPD and soil type and increases with the SWC (*θ*); larger SWC levels give rise to greater water loss (Garratt 1992). During some management period *i*, the SWC changes over time as described by Equation 1:

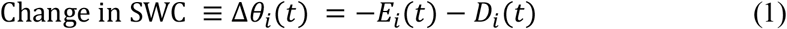

where Δ*θ*_*i*_(*t*) is the change in the SWC per unit of time (e.g., mmol m^-2^ day^-1^). Notice the negative effect of the plant’s own transpiration, *E*_*i*_(*t*), on the SWC.

Following ample evidence (see Letey and Dinar 1986, Shani et al. 2004, Shani et al. 2007, and the references that they cite), we assume that relative biomass growth (i.e., percent change in biomass) at any point in time is proportional to the transpiration rate per unit of biomass, *E*_*i*_(*t*), with the water-use efficiency (*WUE*_*i*_, m^2^ mmol^-1^) as the coefficient of proportionality:

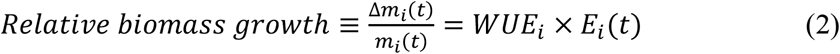

where *m*_*i*_(*t*) is the biomass at time *t* during management period *i*, Δ*m*_*i*_(*t*) is the change in biomass from time *t* − 1 until *t*, and the coefficient *WUE*_*i*_ may change with the biomass (see Appendix A). Due to physiological and anatomical mechanisms that limit the plant’s hydraulic and gas conductance, *E*_i_(*t*) cannot exceed a maximal level, *Ē*, while survival requires a minimal transpiration rate, *<u>E</u>* (i.e., the plant will dry out and die if transpiration drops below *<u>E</u>*). The maximal transpiration rate *Ē* depends on VPD and soil structure and increases as SWC increases (see Fig. 4). The gray area depicted in Fig. 4 represents the set of feasible transpiration rates *E* for a given SWC level *θ*, where the latter is bounded by the field capacity 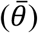 and the wilting SWC, *<u>θ</u>*. At a SWC below *<u>θ</u>*, the plant is unable to extract (transpire) *<u>E</u>* and subsequently dries up.

**Fig 4.**
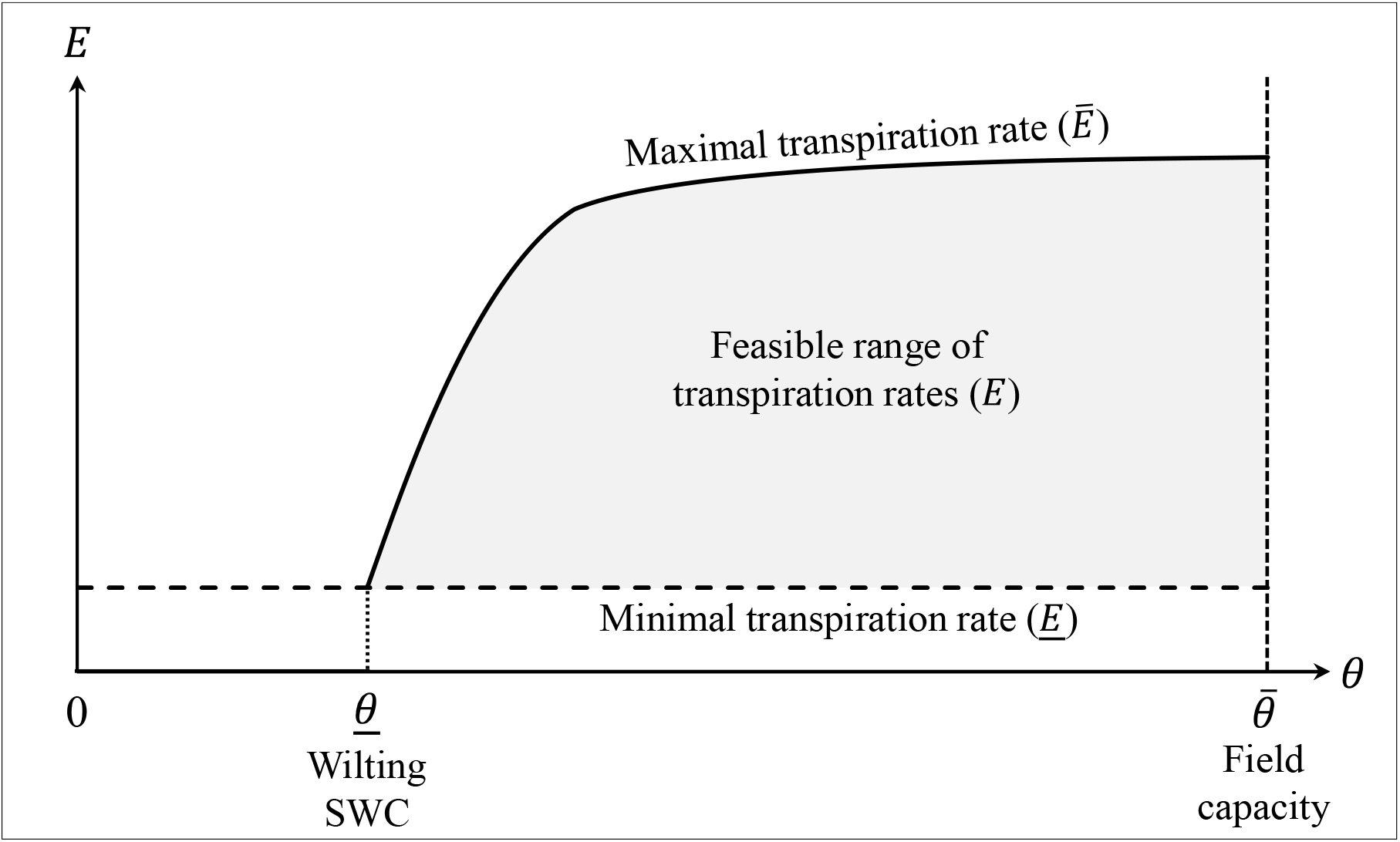
Schematic curve representing the dependence of the upper transpiration bound *Ē* on the level of SWC under constant VPD. The gray area marks the range of feasible transpiration levels (*E*) for any SWC (*θ*) level, where the latter ranges between the field capacity 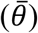 and the wilting SWC level below which the plant desiccates (*<u>θ</u>*), as it cannot transpire the minimal rate required for survival (*<u>E</u>*).

If the SWC reaches the wilting level *<u>θ</u>* before either the next rain event occurs or maturity has been reached, the plant dries up and dies. If the next rain event occurs before the SWC reaches the wilting level, a new management period begins with a replenished SWC. If the plant reaches maturity, the objective has been reached and transpiration is terminated. Because the time at which the next rain event will occur is uncertain, desiccation is also uncertain and depends on the pattern of the transpiration policy: Higher transpiration rates speed up the decline in the SWC (cf. Eq. 1), thereby increasing the likelihood of desiccation (if the next rain event is delayed); whereas low transpiration rates lower the risk of desiccation by pushing off the point at which the SWC will reach the wilting level *<u>θ</u>*. On the other hand, higher transpiration rates give rise to greater biomass growth (cf. Eq. 2) and hence to greater biomass at maturity, if maturity is ever reached. Thus, wild plants face a transpiration dilemma: A higher transpiration rate gives rise to greater biomass growth and a greater risk of desiccation; whereas a lower transpiration rate entails lower biomass growth and a lower risk of desiccation.

The simple fact that a wild plant flourishes in a specific natural habitat implies that, through evolutionary selection processes, it has adapted successfully to that habitat and employs the optimal transpiration policy for that habitat, which maximizes its reproductive prospects (otherwise, it would have been driven out by better adapted varieties or plants). Associating reproductive prospects with expected biomass at maturity, where the expectation accounts for desiccation risks, and identifying the optimal transpiration policy as the one that maximizes expected biomass at maturity resolves the transpiration dilemma. In the following section, we present the optimal transpiration policy in simple terms (a detailed derivation is presented in Appendix A), demonstrate that the optimal policy resembles the observed biphasic transpiration patterns depicted in Fig. 2, and characterize the link between the optimal transpiration policy and the time of maturity.

### Optimal Transpiration Policy

As explained above, each rain event initiates a new management period. The initial SWC at the beginning of Management Period *i, θ*_*i*_(0), depends on the soil type, the intensity of the rain event that initiated the management period and, possibly, the SWC at the end of the previous management period. Given *θ*_*i*_(0), the time until the SWC trajectory *θ*_*i*_(*t*) reaches the wilting level *<u>θ</u>* (if the next rainfall event does not occur beforehand) can be calculated for any feasible transpiration policy *E*_*i*_(*t*) ∈ [*<u>E</u>, Ē*]. In particular, let 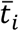and *<u>t</u>*_*i*_ represent the time *θ*_*i*_(*t*) that it takes for the plant to reach the wilting level *<u>θ</u>* under the maximal transpiration condition [i.e., *E*_*i*_(*t*) = *Ē*throughout] and the minimal transpiration condition [i.e., *E*_*i*_(*t*) = *<u>E</u>* throughout]. Clearly 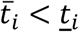(higher transpiration implies a faster decline in SWC and a more rapid arrival at the wilting level *<u>θ</u>*), meaning that survival is guaranteed until 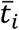, but no later than *<u>t</u>*_*i*_. Fig. 5 depicts schematic trajectories of *E*_*i*_(*t*) and *θ*_*i*_(*t*) during Management Period *i, i*n wh*i*ch the maximal and minimal transpiration behaviors are continuously employed, termed *productive mode* and *survival mode*, respectively. It also shows a *switching-mode* policy, in which transpiration begins at the maximal rate *Ē* and switches to the minimal rate *<u>E</u>* at some time 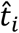 before 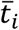; the associated SWC process changes trends at the switching time 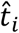 and reaches the witling level at time 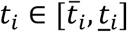.

**Fig 5.**
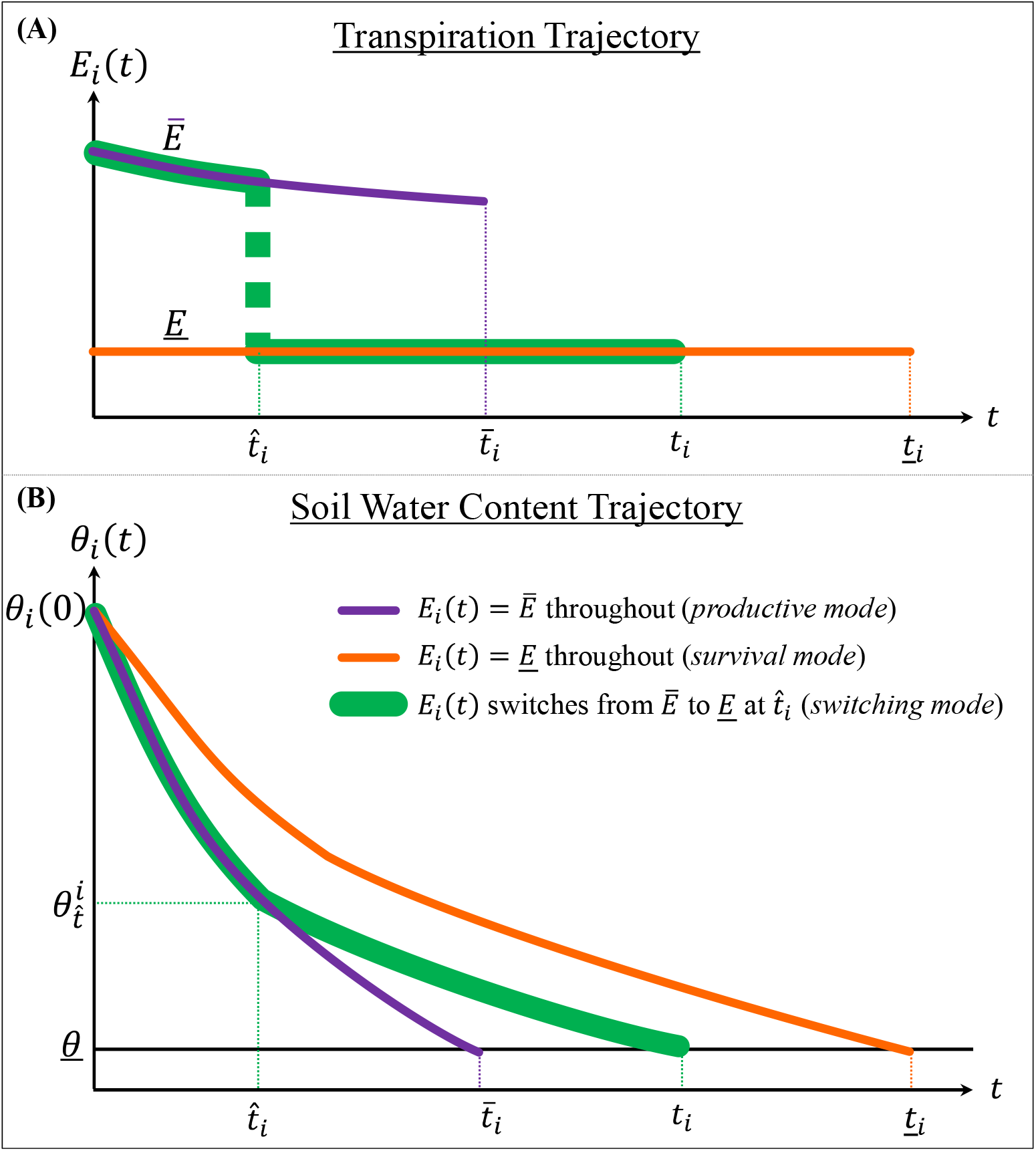
Schematic trajectories of (A) transpiration and (B) SWC during some Management Period *i*, under three transpiration policies: *E*_*i*_(*t*) = *Ē* throughout (*productive mode*), *E*_*i*_(*t*) = *<u>E</u>* throughout (*survival mode*), and *E*_*i*_(*t*) switches from *Ē* to *<u>E</u>* at time 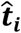 (*switching mode*). Under the productive-mode policy, the SWC path *θ*_*i*_(*t*) declines at the fastest rate and reaches the wilting level *<u>θ</u>* at 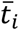(provided the next rainfall event or maturity do not occur beforehand). In survival mode, the SWC trajectory declines at the lowest rate and reaches the wilting level at *<u>t</u>*_*i*_. In the switching mode, the SWC declines at the fastest rate until 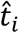 and then switches to the slowest rate, reaching the wilting level at time *t*_*i*_ between 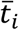 and *<u>t</u>*_*i*_. Notice that *Ē* (A) declines over time due to the declining SWC (recall Fig. 4).

This leads us to Property 1 (see proof in Appendix A).

#### Property 1

*The optimal transpiration policy during a management period admits the switching-mode policy:*

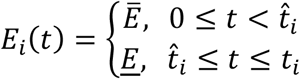

Smoothing out the effects of VPD fluctuations, the trajectories *E*_*i*_(*t*) and *θ*_*i*_(*t*) in the *switching-mode* policy depicted in Fig. 5 (green curves) resemble their counterparts depicted in Fig. 2 [i.e., *E*_*i*_(*t*) = *Ē* and *E*_*i*_(*t*) = *<u>E</u>* during Phase I and II, respectively].

Through switching-mode policy, the maximal transpiration *Ē* is applied first for two main reasons: drainage and uncertain rainfall. Let’s first consider drainage. According to Eq. 1,the management-period’s initial SWC [*θ*_*i*_(0)] is allocated during that period between transpiration and drainage, where both are positively related to SWC. Therefore, for any management period that does not end with desiccation, a switching-mode policy that begins with *Ē* allocates a larger share of *θ*_*i*_(0) to transpiration than switching-mode policy that begins with *E*, implying that the former generates greater biomass growth.

Uncertain rainfall induces a preference for the short term over the long term. To see this, consider the transpiration decision at the beginning of a management period. If the next rainfall event occurs early on, it would certainly be desirable to transpire at the maximal rate early on; whereas if the next rainfall event is delayed, it is still possible to switch to the minimal transpiration and lower the desiccation risk. The uncertain rainfall induces a form of time discounting that causes the plant to behave as if it prefers the short term over the long term (see further discussion in Appendix A).

Property 1 limits the choice of the optimal transpiration policy during any management period to one of the three modes depicted in Fig. 5, in which the cases 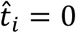 and 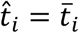 are corner solutions that correspond to the *survival mode* and *productive mode*, respectively. The chosen mode during some Management Period *i* yields the greatest expected biomass growth, given the initial SWC [*θ*_*i*_(0)] and the remaining time to maturity at the onset of the management period, 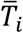, which is the time to maturity from germination, 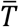, minus the time from germination until the rain event that initiated Management Period *i*. The biomass expectation weighs in the survival (no desiccation) probability; the latter depends on the stochastic nature of the rain events (particularly their frequency and intensity; see Appendix A), drainage and the pattern of the transpiration policy. The specific structure of the three modes of transpiration enables the straightforward calculation of the expected increase in biomass in each mode (see derivation in Appendix A) and, therefore, also the choice of the optimal transpiration mode for each management period over the course of the growing season.

The optimal transpiration policy throughout the growing season evolves along the succession of Management Periods *i* = 1,2, …, each of which is initiated by a rain event.

The management periods differ in two important respects: the initial SWC *θ*_*i*_(0) and the time to maturity 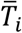. As 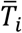decreases as Management Period *i* falls later in the growing season, it gives rise to three types of management periods:

**Type 1:**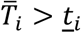, that is, survival until maturity cannot be guaranteed even under minimal transpiration throughout the management period.

**Type 2:**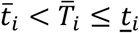, that is, survival until maturity is feasible.

**Type 3:**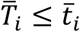, that is, survival until maturity is guaranteed regardless of the pattern of transpiration.

Type 1 management periods fall early in the growing season, when the time to maturity 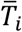is large enough so that, even with minimal transpiration (survival mode), reaching maturity depends on the next rain event occurring early enough – before *<u>t</u>*_*i*_. Type 2 management periods fall later in the growing season, when the time to maturity 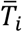is smaller than *<u>t</u>*_*i*_, but larger than 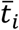, so reaching maturity is feasible. Type 3 management periods occur close enough to the end of the growing season for 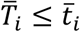, so that survival until maturity is guaranteed even with maximal transpiration throughout. Without desiccation risk, the expected biomass growth is the same as the actual biomass growth, and the latter increases with transpiration (cf. Equation 2). This brings us to Property 2:

#### Property 2

*For Type 3 management periods* (with 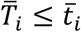), *the productive mode policy (maximal transpiration throughout) is optimal*.

It turns out that for Type 2 management periods, the optimal transpiration trajectory is either the productive mode or the switching mode. The employed mode is the one that yields the higher expected biomass at the end of the management period, as proven in Appendix A.

#### Property 3

*The optimal transpiration during Type 2 management periods* (with 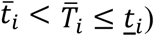) *begins at the maximal rate Ē and lasts until either (i) desiccation occurs at* 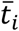*or (ii)* 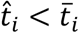, *at which time the plant switches to the minimal transpiration <u>E</u>, where* 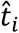 *is chosen such that the SWC process θ*_*i*_(*t*) *will reach the wilting level <u>θ</u> exactly at* 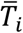*(maturity), i*.*e*.,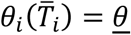.

Property 3(ii) implies that, in the *switching mode* policy, the plant transpires at the maximal rate *Ē* for the longest duration for which reaching maturity is guaranteed by switching to the minimal rate *<u>E</u>*. This optimization condition constitutes the link between the transpiration policy of an annual plant and its maturity time.

The optimal transpiration policy for Type 1 management periods (initiated by rain events early in the growing season) remains to be characterized.

#### Property 4

*For Type 1 management periods (with* 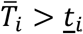*), the optimal transpiration trajectory is the productive mode.*

Property 4 implies that if reaching maturity (survival) cannot be guaranteed even under the (most conservative) survival-mode policy, it is optimal to take the highest (desiccation) risk (by transpiring at the maximal rate) and if the next rain event occurs before desiccation, the plant will enter the next management period with greater biomass. (It turns out that the transpiration pattern described in Property 4 depends on the relationship between the frequency of rainfall events and the minimal biomass accumulated during the period in which the plant acts in survival mode; see discussions in Appendices A and B).

### Natural Selection of Physiological Traits

Our analysis so far has focused on the inner circle of Fig. 1, characterizing the optimal transpiration policy given both the environmental conditions and physiological traits. We defined four key physiological traits: *Ē* = the maximal transpiration rate, *<u>E</u>* = the minimal transpiration rate below which the plant desiccates, *WUE* = water use efficiency (cf. Eq. 2) and 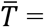the length of the growing season from germination to maturity. We refer to this set of physiological traits as 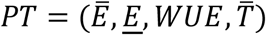. The environmental conditions include soil type, VPD and the frequency and intensity of rainfall events; we refer to the set of environmental conditions as *EV*. Given *PT* and *EV*, the optimal pattern of transpiration throughout the growing season was characterized above (“Optimal Transpiration Policy”). The outcome of this optimal pattern is the expected biomass at maturity, referred to as 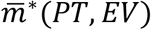.

In this section, we examine the natural selection of the physiological traits *PT* (the middle circle of Fig. 1) in light of the patterns of the optimal transpiration policy, given the environmental conditions *EV* and subject to biological constraints (outer circle of Fig. 1). The underlying hypothesis is that the observed traits have been adjusted (via natural selection) to the prevailing environmental conditions, in order to improve the performance of the optimal transpiration policy. That is, the observed traits maximize 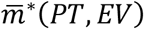 subject to biological constraints, given the environmental conditions *EV*. The outcome of this selection process is referred to as *PT*^∗^(*EV*).

Our analysis integrated the optimal transpiration principles described above (“Optimal Transpiration Policy”) with empirical measures of physiological traits and environmental conditions. To elicit measures of the physiological traits *WUE, Ē* and *Ē*, we developed an econometric procedure to isolate the impact of the plant’s mechanisms (e.g., stomatal apertures) on its transpiration from the effects of VPD and SWC (see Appendix B). We applied this procedure to the wild-barley accessions sampled from the aforementioned five natural sites (i.e., Meron, Arbel, Oren, Guvrin and Yeruham) using experimental data reported by Galkin et al. (2018).

Suppose that nature could unlimitedly select plants’ physiological traits and let 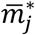represent the direct (partial) effect of a small (marginal) change in Physiological Trait *j* on 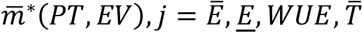. Based on the optimal transpiration policy characterized above, these partial effects satisfy:

#### Property 5

(i) 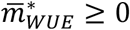(ii) 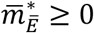and 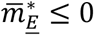

**Proof:** (i) Suppose that, ceteris paribus, *WUE* is increased slightly to *WUE* + Δ. Then, the optimal transpiration policy under *WUE* is feasible and yields a larger biomass under *WUE* + Δ, implying that 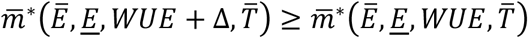, verifying (i). To verify (ii), suppose that *Ē* is increased slightly to *Ē* + Δ. Then, the optimal policy under *E* is feasible under *Ē* + Δ, implying that 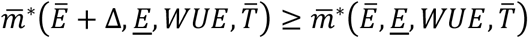. Finally, to verify (iii) suppose that *<u>E</u>* is increased slightly to *<u>E</u>* + Δ. Then, the optimal policy under *<u>E</u>* + Δ is feasible under *<u>E</u>* + Δ, implying that 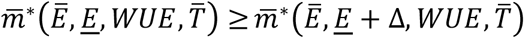. □

Property 5(i) is obvious: greater biomass growth per unit of transpiration can only improve yield (cf. Eq. 2). Property 5(ii) follows from the observation that higher maximal transpiration rates allow for the decreased loss of water to drainage (cf. Eq. 1). Property 5(iii) reflects the simple fact that lower minimal transpiration rates reduce the risk of desiccation when the timing of the next rain event is uncertain. Notice that, in view of Property 3(ii), the sign of 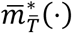is indeterminate. That is, it can be negative, positive or zero, depending on the other traits and environmental conditions (particularly rainfall and drainage patterns).

Property 5 implies that an ideotypic annual plant is characterized by *WUE* = ∞, *Ē* = ∞ and *<u>E</u>* = 0 — a combination that enables immediate transpiration of all of the soil water accessible to the plant following any rainfall event together with perfect transformation of the transpired water into biomass, and then survival until the next rainfall event or maturity regardless of their timing. We claim that such an ideotype does not exist because of biological constraints that impose tradeoffs between these three physiological traits.

Based on our empirical measures of the *WUE, Ē* and *<u>E</u>* of the five wild barley accessions, we identified two biological constraints (see Fig. 6):

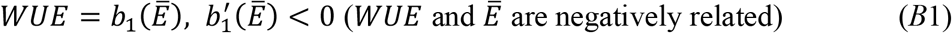

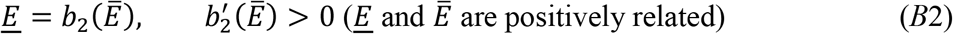

The *WUE*-*Ē* constraint (*B*1; Fig. 6A) stems from the saturation of net CO_2_ assimilation at high transpiration rates due to limitations on the regeneration of ribulose-1,5-bisphosphate (see Yoo et al., 2009). A possible explanation for the *<u>E</u>*-*Ē* constraint (*B*2; Fig. 6B) is that a higher level of *Ē* requires a larger number of stomata per unit of leaf area and/or wider stomatal apertures, both of which entail greater water loss when the plant is in survival mode (i.e., larger *<u>E</u>*). The *<u>E</u>*-*Ē* constraint may be attributed to the multi-functionality of stomata, which should enable fast and slow transpiration rates when the plant is in its productive and survival modes, respectively. That is, this limitation reflects the fact that “a given phenotype cannot be optimal at all tasks” (Shoval et al. 2012, p. 1157).

**Fig 6.**
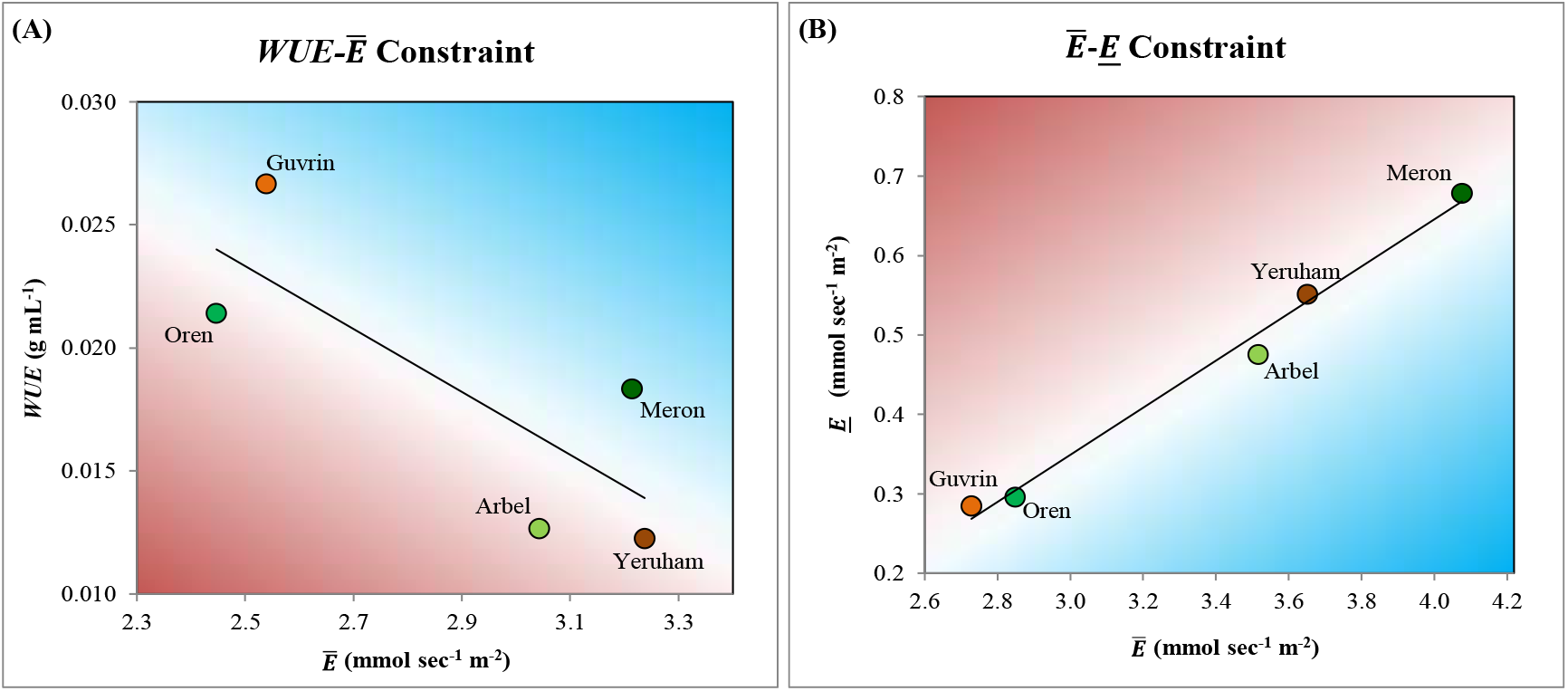
*WUE, Ē* and *E*. **(A)** *WUE* plotted against *Ē* (both measured under non-stressed conditions), with the linear regression line reflecting the *WUE*-*Ē* constraint (*B*1). **(B)** *<u>E</u>* plotted against *Ē* (measured under water-deficit conditions) with the linear regression line representing the *<u>E</u>*-*Ē* constraint (*B*2). The blue (red) areas represent trait combinations that allow for greater (smaller) biomass at maturity.

Constraint *B*1 limits the possibility of increasing both *WUE* and *Ē* (i.e., selecting combinations that fall within the blue area of Fig. 6A), in order to accelerate biomass accumulation (cf. Eq. 2). Constraint *B*2 limits the possibility of *<u>E</u>*-*Ē* combinations that fall within the blue area in Fig. 6B. Consequently, increasing *Ē* to enhance growth and reduce the amount of water lost to drainage when the plant is in productive mode comes at the expense of a larger *<u>E</u>*, which accelerates the decline in the SWC when the plant is in survival mode, which increases the risk of desiccation.

The direct effects in Property 5 ignore the biological constraints *B*1 and *B*2. Incorporating the latter requires accounting for the total effect on 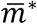 of each trait [i.e., the direct (partial) effects plus the indirect effects induced by the biological constraints]. The environmentally selected optimal set of traits in a given environment, *PT*^∗^(*EV*), maximizes the expected biomass at maturity 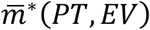subject to the biological constraints (*B*1 and *B*2).

We now use Property 5 combined with the two biological constraints to explain the observed physiological traits [i.e., *PT*^∗^(*EV*)] in the five habitats from which our wild-barley accessions were sampled (Appendix B). The habitats vary widely in their environmental conditions. The northern-most site, Meron, is relatively rainy and cold; whereas the southern-most site, Yeruham, is arid, with infrequent precipitation that is relatively non-intense, as well as sandy-loess soil with a high bulk density, which leads to low water retention and fast drainage (Tian et al. 2018), the rapid drying out of soil pores (Shokri and Or, 2011) and a soil crust that leads to a low level of rainwater infiltration (Eldridge et al. 2000). The habitats of Oren, Arbel and Guvrin are characterized by intermediate conditions.

In Fig. 7A, we show how a larger precipitation level gives rise to a larger 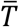, consistent with previous findings (e.g., Aronson et al. 1992). This is because more frequent and intense rain events reduce the risk of desiccation and enable the utilization of potentially more rainfall events for biomass accumulation throughout the growing season. In Meron, the relatively low desiccation risk associated with intense and frequent rain events and the low VPD at that site (Fig. 7D) enable relatively high 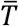(Fig. 7A) and *Ē* (Fig. 7B); the latter (recalling *B*2) entails relatively higher *<u>E</u>* (Fig. 6B). In contrast, the paucity of rain events and low rainwater infiltration and soil water retention in Yeruham give rise to a smaller 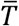. To reach a critical biomass at maturity and cope with the rapid evaporation (i.e., drainage) associated with the bulky soil (Fig. 7C), growth in Yeruham must be rapid, explaining the large *Ē* for that site, relative to the intermediate habitats (Fig. 7B).

**Fig 7.**
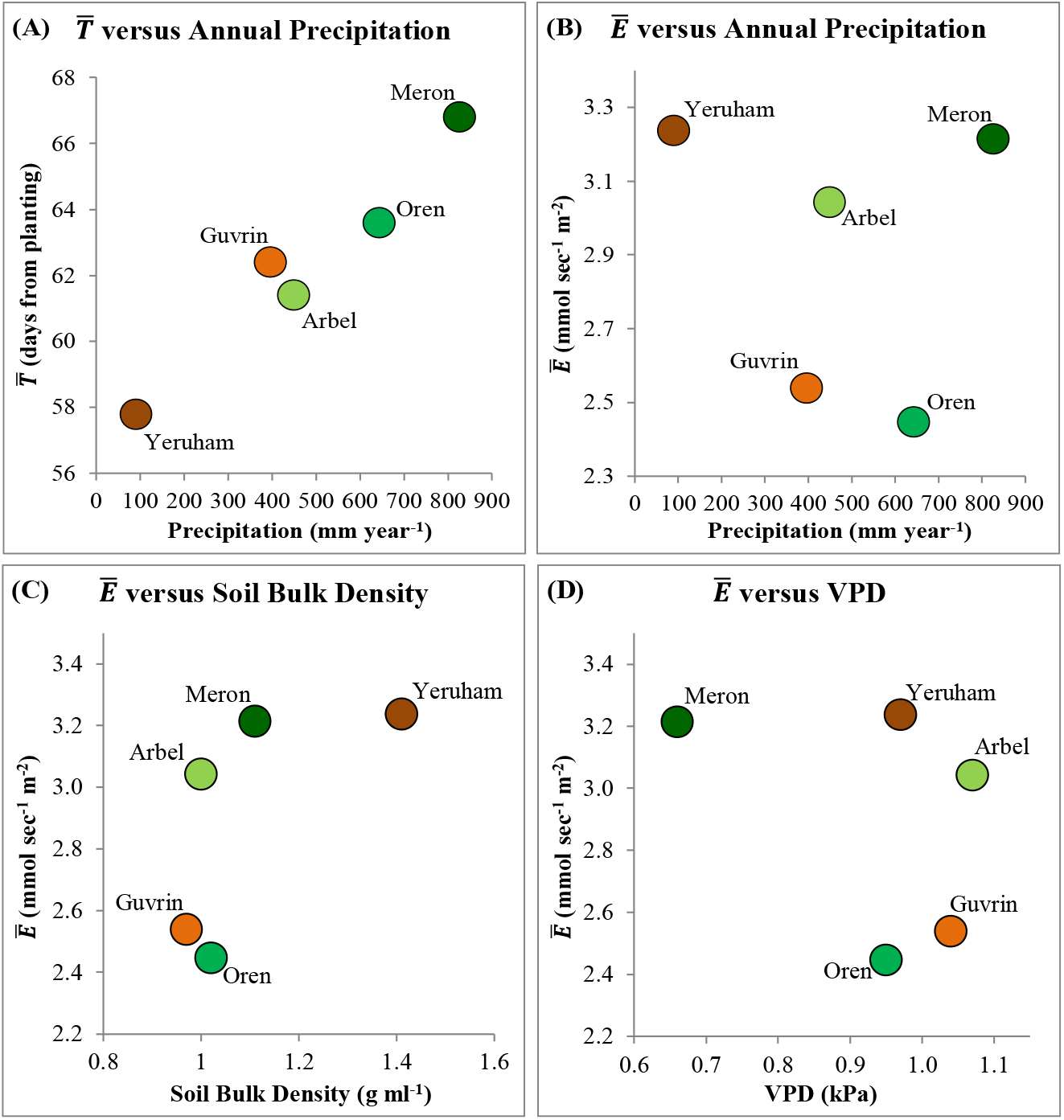
Measured 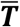and *Ē* plotted against annual precipitation (A and B, respectively) and measured *Ē* plotted against soil bulk density and VPD (C and D, respectively).

Our empirical observations indicate that all traits are changeable in the process of adaptation to different environments, but 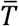seems to be more adaptable than the other examined traits, as indicated by its strong correlation with precipitation (Fig. 7A). This finding is consistent with genetic analyses of wild annual plants, which identified “the flowering-time quantitative trait loci as a driver of adaptive population divergence” (Verhoeven et al. 2008, p. 3416).

Similarly, Richards et al. (2010) pointed to flowering time as the most important physiological trait for the adaptation of commercial cereal crops to Australia’s dry environments. Recalling Property 3ii, 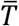is linked to the other traits only via the patterns of the optimal transpiration policy; whereas *WUE, Ē* and *<u>E</u>* are also linked to one another through the biological constraints (*B*1 and *B*2). Thus, 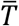is more flexible than the other traits and, therefore, responds more strongly to changing environmental conditions.

### Breeding Implications

When the goal of a breeding program is to increase the yield of a specific rain-fed annual crop in a particular environment, the traits of wild ancestors of that crop in a similar environment can provide useful information, because those traits were naturally selected to maximize reproductive fitness (or expected biomass at maturity) in that environment. As an illustration, suppose that breeders produce a set of germplasm candidate plants, select a genetically-related local wild plant as a benchmark and then measure the three biologically constrained traits *WUE, Ē* and *<u>E</u>* for all plants using the pre-field procedures described in Appendix B. The gray ellipsoids in Fig. 8 present possible distributions of trait combinations of the germplasm candidate plants plotted in the *WUE*-*Ē* plane (Fig. 8A) and the *<u>E</u>*-*Ē* plane (Fig. 8B). The ellipsoids’ inclinations follow from the biological constraints discussed above (Fig. 6). The yellow dots represent the evolutionarily selected trait combinations of the local wild benchmark [i.e., *PT*^∗^(*EV*)]. The polygons marked *a* and *b* in Fig. 8 represent a range of trait combinations that are anchored around the naturally selected combination *PT*^∗^(*EV*), and also induce greater expected biomass at maturity than *PT*^∗^(*EV*), as implied by the partial trait effects 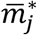discussed above (“Natural Selection of Physiological Traits”). To select the best candidates for the next phases of the breeding program, breeders should choose candidates whose trait combinations fall within both polygons. This type of pre-field screening is likely to make the breeding of rain-fed annual crops for a given environment quicker and cheaper.

**Fig 8.**
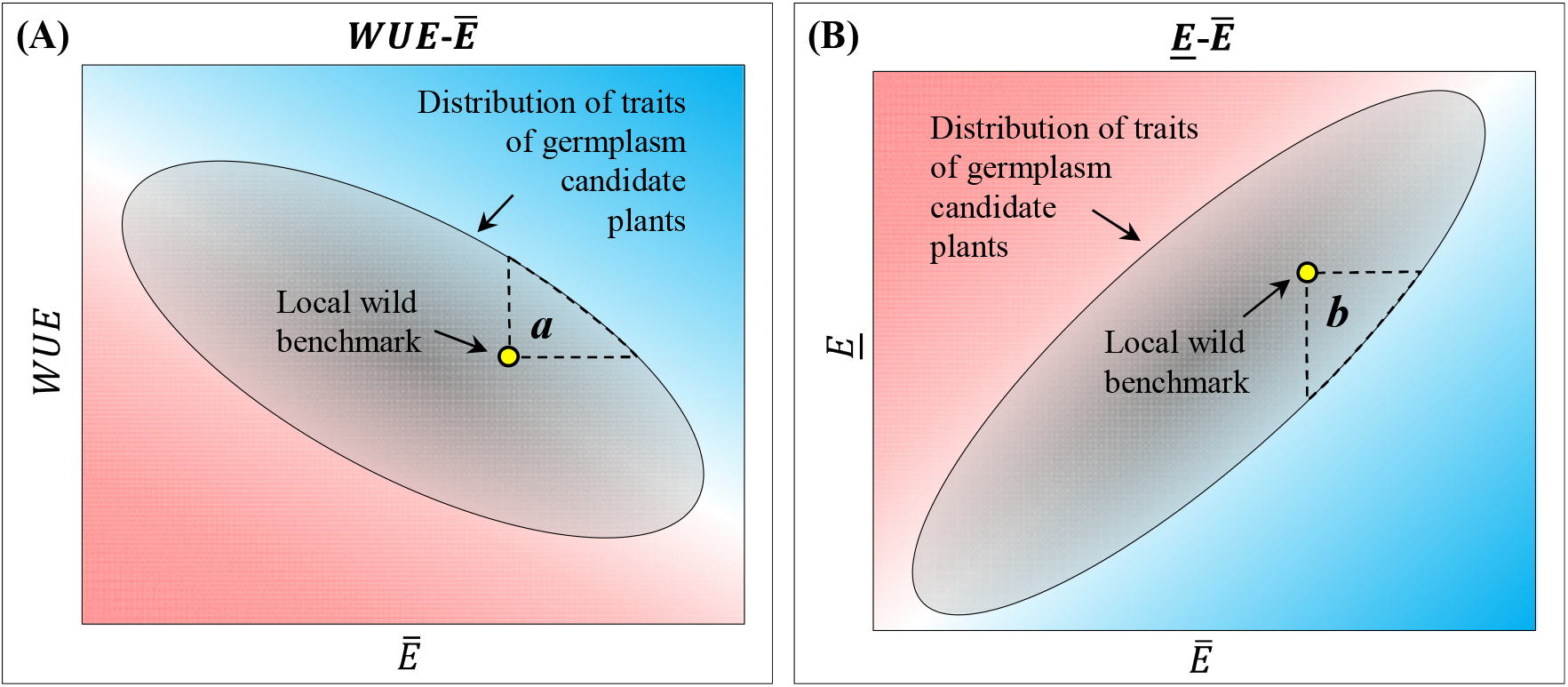
Schematic illustration of criteria for the selection of candidate annual crop lines for rain-fed production in a specific environment based on their combined *WUE, Ē* and *<u>E</u>* traits [whose distributions are represented by the gray ellipsoids in panels (A) and (B)] relative to those of a local wild genotype (yellow dots). Candidates whose trait combinations fall within the dashed-line polygons (near the yellow dots, toward the blue areas) denoted *a* and *b* in panels (A) and (B), respectively, are likely to have larger expected yield in the wild genotype’s environment relative to candidates whose trait combinations fall out of the polygons.

### Concluding Remarks

This study characterizes the optimal transpiration policy of wild annual plants in environments with stochastic rainfall, identifies the key physiological traits governing that policy and their desired levels from the perspective of maximal fitness or expected reproduction, and deduces from experimental output the presence of biological constraints that impose tradeoffs between these traits. Due to these biological constraints, ideotypic plants do not exist and the plants in each environment are characterized by a specific optimal combination of traits. This theory, supplemented with practical trait-quantification methods (Appendix B), constitutes a comprehensive analytical framework for both breeding and research. It may help to explain differences in the geographical spread of different species (e.g., the spread of wild barley versus that of wild emmer wheat; see Peleg et al. 2005) and to characterize the mechanisms and genetics underlying the biological constraints that limit the development of highly productive annual crops for rainfed agriculture in stochastic-rainfall environments.

## List of supplementary materials

Appendix A - Proofs of optimal transpiration properties.

Appendix B – Empirical analyses.

## Funding

This research was partially funded by the Center for Research in Agriculture, Environmental Quality and Natural Resources at the Hebrew University of Jerusalem, and by the Israeli Ministry of Agriculture and Rural Development (grant no. 039-9274).

## Author contributions

IK and MM defined the research question; YT conducted the theoretical analyses; MM provided experimental data and biological insights; IK conducted the empirical analyses, and wrote the manuscript with the help of YT and MM.

## Competing interests

The authors declare no competing interests.

## Data and materials availability

The experimental data used in this study were previously used by studies published in scientific articles, as specified throughout the manuscript. The data and detailed information on the empirical procedures and results are available from the corresponding author upon request.

## Notes

### Competing Interest Statement

The authors have declared no competing interest.

## References

Aronson, J., Kigel, J., Shmida, A., & Klein, J. 1992. Adaptive phenology of desert and Mediterranean populations of annual plants grown with and without water stress. Oecologia, 89(1), 17–26.

Blum, A. (2009). Effective use of water (EUW) and not water-use efficiency (WUE) is the target of crop yield improvement under drought stress. Field Crops Research, 112(2-3), 119–123.

Cowan, I. R., & Farquhar, G. D. (1977). Stomatal function in relation to leaf metabolism and environment. In Symposia of the Society for Experimental Biology (Vol. 31, p. 471).

Dalal, A., Attia, Z., & Moshelion, M. (2017). To produce or to survive: How plastic is your crop stress physiology?. Frontiers in Plant Science, 8, 2067.

Eldridge, D. J., Zaady, E., & Shachak, M. (2000). Infiltration through three contrasting biological soil crusts in patterned landscapes in the Negev, Israel. Catena, 40(3), 323–336.

Food and Agriculture Organization (2017). The future of food and agriculture – trends and challenges, Rome: FAO.

Galkin, E., Dalal, A., Evenko, A., Fridman, E., Kan, I., Wallach, R. & Moshelion, M. (2018). Risk-management strategies and transpiration rates of wild barley in uncertain environments. Physiologia Plantarum, 164(4), 412–428.

Garratt, J. R. (1992). The atmospheric boundary layer. Cambridge Univ. Press, Cambridge, 316.

Graff, G., Hochman, G., & Zilberman, D. (2013). The research, development, commercialization, and adoption of drought and stress-tolerant crops, in Crop Improvement under Adverse Conditions, eds N. Tuteja and S. S. Gill (New York, NY: Springer), 1–33. doi: 10.1007/978-1-4614-4633-0_1.

Großkinsky, D. K., Svensgaard, J., Christensen, S., & Roitsch, T. (2015). Plant phenomics and the need for physiological phenotyping across scales to narrow the genotype-to-phenotype knowledge gap. Journal of Experimental Botany, 66(18), 5429–5440.

Hübner, S., Bdolach, E., Ein-Gedy, S., Schmid, K. J., Korol, A., & Fridman, E. (2013). Phenotypic landscapes: phenological patterns in wild and cultivated barley. Journal of Evolutionary Biology, 26(1), 163–174.

Hübner, S., Hoffken, M., Oren, E., Haseneyer, G., Stein, N., Graner, A., Schmid, K. & Fridman, E. (2009). Strong correlation of wild barley (Hordeum spontaneum) population structure with temperature and precipitation variation. Molecular Ecology, 18, 1523–1536.

Letey, J., & Dinar, A. (1986). Simulated crop-water production functions for several crops when irrigated with saline waters. Hilgardia, 54(1), 1–32.

Lobell, D. B., Roberts, M. J., Schlenker, W., Braun, N., Little, B. B., Rejesus, R. M., & Hammer, G. L. (2014). Greater sensitivity to drought accompanies maize yield increase in the US Midwest. Science, 344(6183), 516–519.

Lu, Y., Duursma, R. A., & Medlyn, B. E. (2016). Optimal stomatal behaviour under stochastic rainfall. Journal of Theoretical Biology, 394, 160–171.

Mäkelä, A., Berninger, F., & Hari, P. (1996). Optimal control of gas exchange during drought: theoretical analysis. Annals of Botany, 77(5), 461–468.

Mencuccini, M., Manzoni, S., & Christoffersen, B. (2019). Modelling water fluxes in plants: from tissues to biosphere. New Phytologist, 222(3), 1207–1222.

Merchuk-Ovnat, L., Silberman, R., Laiba, E., Maurer, A., Pillen, K., Faigenboim, A., & Fridman, E. (2018). Genome scan identifies flowering-independent effects of barley HsDry2. 2 locus on yield traits under water deficit. Journal of Experimental Botany, 69(7), 1765–1779.

Mickelbart, M. V., Hasegawa, P. M., & Bailey-Serres, J. (2015). Genetic mechanisms of abiotic stress tolerance that translate to crop yield stability. Nature Reviews Genetics, 16(4), 237–251.

Millet, E. J., Kruijer, W., Coupel-Ledru, A., Prado, S. A., Cabrera-Bosquet, L., Lacube, S., Charcosset, A., Welcker, C., van Eeuwijk, F. & Tardieu, F. (2019). Genomic prediction of maize yield across European environmental conditions. Nature Genetics, 51(6), 952–956.

Moshelion, M. & Altman, A. (2015). Current challenges and future perspectives of plant and agricultural biotechnology. Trends in Biotechnology, 33, 337–342.

Moshelion, M., Halperin, O., Wallach, R., Oren, R. A. M., & Way, D. A. (2015). Role of aquaporins in determining transpiration and photosynthesis in water-stressed plants: crop water-use efficiency, growth and yield. Plant, Cell and Environment, 38(9), 1785–1793.

Negin, B., & Moshelion, M. (2017). The advantages of functional phenotyping in pre-field screening for drought-tolerant crops. Functional Plant Biology, 44(1): 107–118.

Peleg, Z., Fahima, T., Abbo, S., Krugman, T., Nevo, E., Yakir, D., & Saranga, Y. (2005). Genetic diversity for drought resistance in wild emmer wheat and its ecogeographical associations. Plant, Cell & Environment, 28(2), 176–191.

Richards, R. A., Rebetzke, G. J., Watt, M., Condon, A. T., Spielmeyer, W., & Dolferus, R. (2010). Breeding for improved water productivity in temperate cereals: phenotyping, quantitative trait loci, markers and the selection environment. Functional Plant Biology, 37(2), 85–97.

Shani, U., Ben-Gal, A., Tripler, E., & Dudley, L. M. (2007). Plant response to the soil environment: An analytical model integrating yield, water, soil type, and salinity. Water Resources Research, 43(8).

Shani, U., Y. Tsur and A. Zemel (2004) Optimal dynamic irrigation schemes, Optimal Control Applications and Methods 25: 91–106.

Shokri, N. and Or, D. (2011). What determines drying rates at the onset of diffusion controlled stage-2 evaporation from porous media?. Water Resources Research, 47(9).

Shoval, O., Sheftel, H., Shinar, G., Hart, Y., Ramote, O., Mayo, A., Dekel, E., Kavanagh, K. & Alon, U. (2012). Evolutionary trade-offs, Pareto optimality, and the geometry of phenotype space. Science, 336(6085), 1157–1160.

Skirycz, A., Vandenbroucke, K., Clauw, P., Maleux, K., De Meyer, B., Dhondt, S., Pucci, A., Gonzalez, N., Hoeberichts, F., Tognetti, V. B., Galbiati, M., Tonelli, C., Van Breusegem, F., Vuylsteke, M., & Inzé, D. (2011). Survival and growth of Arabidopsis plants given limited water are not equal. Nature Biotechnology, 29(3), 212–214.

Sultan, S. E. (2000). Phenotypic plasticity for plant development, function and life history. Trends in Plant Science, 5(12), 537–542.

Takeda, S., & Matsuoka, M. (2008). Genetic approaches to crop improvement: responding to environmental and population changes. Nature Reviews Genetics, 9(6), 444–457.

Tian, Z., Gao, W., Kool, D., Ren, T., Horton, R., and Heitman, J. L. (2018). Approaches for estimating soil water retention curves at various bulk densities with the extended van Genuchten model. Water Resources Research, 54(8), 5584–5601.

Tilman, D., Balzer, C., Hill, J., & Befort, B. L. (2011). Global food demand and the sustainable intensification of agriculture. Proceedings of the National Academy of Sciences, 108(50), 20260–20264.

Tuberosa, R., Turner, N. C. & Cakir, M. (2014). Two decades of InterDrought conferences: are we bridging the genotype-to-phenotype gap? Journal of Experimental Botany, 65, 6137–6139.

Verhoeven, K. J. F., Poorter, H., Nevo, E., & Biere, A. (2008). Habitat-specific natural selection at a flowering-time QTL is a main driver of local adaptation in two wild barley populations. Molecular Ecology, 17(14), 3416–3424.

Volis, S., Verhoeven, K. J. F., Mendlinger, S., & Ward, D. (2004). Phenotypic selection and regulation of reproduction in different environments in wild barley. Journal of evolutionary biology, 17(5), 1121–1131.

Wani, S.P., Rockström, J., & Oweis, T. Y. (Eds.). (2009). Rainfed agriculture: unlocking the potential (Vol. 7). CABI.

Yoo, C. Y., Pence, H. E., Hasegawa, P. M., & Mickelbart, M. V. (2009). Regulation of transpiration to improve crop water use. Critical Reviews in Plant Science, 28(6), 410–431.

Zhao, C., Liu, B., Piao, S., Wang, X., Lobell, D. B., Huang, Y., Huang, M., Yao, Y., Bassu, S., Ciais, P., Durand, J. L., Elliott, J., Ewert, F., Janssen, I. A., Li, T., Lin, E.,Liu, Q., Martre, P., Müller, C., Peng, S., Peñuelas, J., Ruane, A. C., Wallach, D., Wang, T., Dongha, W., Liu, Z., Zhu, Z., and Asseng, S. (2017). Temperature increase reduces global yields of major crops in four independent estimates. Proceedings of the National Academy of Sciences, 114(35), 9326–9331.

