## Appendix A for "On the biological constraints that limit the productivity of rain-fed annual crops"

Supplementary materials for:
On the Biological Constraints that Limit the Productiveness of Rainfed Annual Crops

Yacov Tsur[†]

### A Proofs of optimal transpiration properties

#### Setting

Rainfall occurs at discrete times $\tau_0$, $\tau_1, \tau_2, \ldots$, that follow a series of Poisson events with a constant mean arrival rate $\lambda$, where $\tau_0 = 0$ represents the germination time (occurs at the onset of a rain event). With $\tilde{T}_i = \tau_i - \tau_{i-1}$ representing the time interval between consecutive rain events, $\tilde{T}_i$, $i = 1, 2, \ldots$, are i.i.d. draws from the distribution

$$Pr\{\tilde{T}_i \leq t\} = 1 - e^{-\lambda t} \tag{A.1}$$

with mean $E\{\tilde{T}_i\} = 1/\lambda$. A rainy region (with frequent rain events) is characterized by a large $\lambda$ and small expected time between rain events. An arid region, with infrequent rain events, has a smaller $\lambda$ and larger expected time ($1/\lambda$) between rain events.

The growing season (from germination to maturity) is of length $\bar{T}$ and progresses along rain events, each marking the beginning of a *management period* that lasts until either the next rain event occurs or the plant reaches maturity or the plant desiccates, whatever is encountered first. The transpiration policy is determined for each management period. Management periods differ with respect to their initial soil water content (SWC), initial biomass and the remaining time to maturity $\bar{T}_i = \bar{T} - \tau_{i-1}$. The initial SWC depends on the intensity of the rain event at the onset of the management period; the initial biomass and time to maturity vary along the growing season (see Figure 3).

[†]Department of Environmental Economics and Management and the Center for Agricultural Economic Research, The Hebrew University of Jerusalem, POB 12, Rehovot 7610001, Israel.

The $i$'th rain event instantly replenishes the SWC at the root zone to the level $\theta_i(0) \leq \bar{\theta}$, where $\bar{\theta}$ represents saturation (field capacity) and variation in $\theta_i(0)$ stems from variation in the intensity and frequency of rainfall events. Until the next rain event, the SWC decreases with per unit biomass transpiration $E_i(t)$ and per unit biomass drainage $D(\theta_i(t))$ according to

$$\dot{\theta}_i(t) = -E_i(t) - D(\theta_i(t)), \tag{A.2}$$

where $D(\cdot)$ is increasing in the SWC and time $t$ is measured from the beginning of management period $i$, i.e., from $\tau_{i-1}$. This process ends when the $i$'th rain event occurs, at which time the SWC is replenished and its dynamics evolves as described in (A.2) for management period $i+1$ and so on.

The plant's biomass during management period $i$, denoted $m_i(t)$, grows according to

$$\dot{m}_i(t) = E_i(t)g(m_i(t)),\ m_i(0) = m(\tau_{i-1}), \tag{A.3}$$

provided $E_i(t) \geq \underline{E} \geq 0$, where $g(m)$ equals water use efficiency (denoted $\alpha$) time biomass and $\underline{E}$ is the minimal transpiration rate needed to avoid desiccation (see Shani et al. 2004). The process $m(t)$ without the subscript $i$ measures biomass from germination, so $m_i(0) = m(\tau_{i-1})$. The plant's transpiration rate is bounded between $\bar{E}(\theta)$ and $\underline{E}$. The wilting SWC level $\underline{\theta}$ is defined by (Figure 4, Section 3)

$$\bar{E}(\underline{\theta}) = \underline{E}. \tag{A.4}$$

If $\theta_i(t)$ decreases below $\underline{\theta}$, the plant can no longer extract the subsistence water dose $\underline{E}$ needed for survival and desiccates. The uncertainty regarding the timing of the next rain event induces an uncertainty on the event that $\theta_i(t)$ decreases below $\underline{\theta}$ and triggers desiccation.

#### The optimal transpiration policy

The optimal transpiration policy evolves along successive management periods until maturity or desiccation, whatever occurs first. At each point of time during the $i$'th management period, the plant faces the following dilemma. Increasing $E_i(t)$ raises biomass growth (Eq. (A.3)) but at the same time accelerates the rate at which $\theta_i(t)$ declines (Eq. (A.2)), thereby increasing the probability that $\theta_i(t)$ will decline below the wilting level $\underline{\theta}$ before the next rain event occurs, triggering desiccation (if maturity has not been reached yet). The optimal transpiration policy resolves this dilemma. While plants (typically) do not solve optimization problems (not even at night when they are not busy transpiring), evolution selects varieties that behave optimally and drives to extinction those that do not.

The optimal transpiration policy during management period $i$ utilizes data observed at $\tau_{i-1}$ – the onset of the $i$'th management period. The first observation is the just replenished SWC level $\theta_i(0)$, which allows calculating the time $\theta_i(t)$ will reach the wilting level $\underline{\theta}$ in the absence of rain events for any feasible transpiration policy. Of particular importance are the times $\bar{t}_i$ and $\underline{t}_i$ at which the SWC process that departs from $\theta_i(0)$ and evolves according to (A.2) will reach $\underline{\theta}$ under the maximal and minimal transpiration policies, respectively. Noting (A.2), these times satisfy (are defined by)

$$\int_0^{\bar{t}_i} \left[\bar{E}(\bar{\theta}_i(t)) + D\left(\bar{\theta}_i(t)\right)\right] dt = \theta_i(0) - \underline{\theta} \tag{A.5a}$$

and

$$\int_0^{\underline{t}_i} \left[\underline{E} + D\left(\underline{\theta}_i(t)\right)\right] dt = \theta_i(0) - \underline{\theta}, \tag{A.5b}$$

where $\bar{\theta}_i(t)$ and $\underline{\theta}_i(t)$ are the SWC processes that depart from $\theta_i(0)$ under maximal transpiration and minimal transpiration, respectively. Obviously, $\bar{t}_i < \underline{t}_i$.

The second useful observation is the time to maturity

$$\bar{T}_i = \bar{T} - \tau_{i-1}, \tag{A.6}$$

which identifies three types of management periods, depending on whether $\bar{T}_i > \underline{t}_i$ (Type 1), $\bar{t}_i < \bar{T}_i \leq \underline{t}_i$ (Type 2), or $\bar{T}_i \leq \bar{t}_i$ (Type 3). Type 1 management periods fall early in the growing season when maturity is faraway and cannot be reached without the help of future rain events. For Type 2 management periods, which fall later in the growing season, reaching maturity without the help of future rain events is feasible but requires transpiring below the maximal rate part of the time. Type 3 management periods fall close enough to maturity and reaching maturity is guaranteed even under maximal transpiration throughout, disregarding future rain events.

Noting (A.3), it is obvious that:

**Proposition 1.** *For Type 3 management periods ($\bar{T}_i \leq \bar{t}_i$), the optimal policy is maximal transpiration until maturity.*

Proposition 1 verifies Property 2.

For Type 1 or Type 2 management periods reaching maturity without the help of rain events is non feasible or requires transpiring below the maximal rate part of the time, respectively. Let $t_i$ denote the time at which $\theta_i(t)$ reaches the wilting level $\underline{\theta}$ in the absence of rain events, i.e., $\theta(t_i) = \underline{\theta}$. Noting (A.2), $t_i$ satisfies

$$\int_0^{t_i} [E_i(t) + D(\theta_i(t))]dt = \theta_i(0) - \underline{\theta}\,, \tag{A.7}$$

where $\theta_i(t)$ is the SWC process that departs from $\theta_i(0)$ and evolves in time according to (A.2) under a feasible $E_i(t) \in [\underline{E}\,, \bar{E}(\theta_i(t))]$ transpiration policy. Notice that $t_i$ depends on the transpiration policy $E_i(t)$ and the restriction $E_i(t) \in [\underline{E}\,, \bar{E}(\theta_i(t))]$ implies, noting (A.5),

$$t_i \in [\bar{t}_i, \underline{t}_i]. \tag{A.8}$$

Clearly, if setting $t_i > \bar{T}_i$ is feasible (e.g., for Type 2 management periods) it cannot be optimal, as it implies the existence of an alternative policy, with $t_i = \bar{T}_i$ and higher transpiration rate at the beginning of the management period, under which reaching maturity is guaranteed yet biomass growth (noting (A.3)) is higher. The biomass at maturity under the latter policy is therefore higher than that under the policy with $t_i > \bar{T}_i$, implying that the latter cannot be optimal. Thus,

**Lemma 1.** *For Type 2 management periods, $t_i \leq \bar{T}_i$ under the optimal policy.*

The choice of $t_i \leq \bar{T}_i$ requires comparing the expected biomass growth under $t_i < \bar{T}_i$ and under $t_i = \bar{T}_i$. The latter eliminates the desiccation risk and, noting (A.3), gives the biomass growth

$$\int_0^{\min(\tilde{T}_i, \bar{T}_i)} E_i(t) g(m_i(t)) dt.$$

Letting $I(\cdot)$ represent the indicator function that obtains the value one or zero when its argument is true or false, respectively, this biomass growth can be expressed as

$$\int_0^{\bar{T}_i} E_i(t) g(m_i(t)) I(\tilde{T}_i > t) dt.$$

Taking expectation with respect to $\tilde{T}_i$, noting (A.1), the expected biomass growth when $t_i = \bar{T}_i$ equals

$$\int_0^{\bar{T}_i} E_i(t) g(m_i(t)) e^{-\lambda t} dt. \tag{A.9}$$

Under $t_i < \bar{T}_i$, desiccation occurs if $\tilde{T}_i > t_i$ (i.e., the next rain event occurs after $t_i$), leading to loss of the entire biomass, hence the biomass growth in this case equals

$$I(\tilde{T}_i \leq t_i) \int_0^{\tilde{T}_i} E_i(t) g(m_i(t)) dt + I(\tilde{T}_i > t_i) \times 0.$$

Expressing the integral above as $\int_0^{t_i} E_i(t)g(m_i(t))I(\tilde{T}_i \geq t)dt$ and taking expectation, the expected biomass growth under $t_i < \bar{T}$ can be expressed as

$$\left(1 - e^{-\lambda t_i}\right) \int_0^{t_i} E_i(t)g(m_i(t))e^{-\lambda t}dt. \tag{A.10}$$

The $e^{-\lambda t}$ term in the integrand is akin to a discount factor. The associated discount rate, $\lambda$, measures the frequency of rain events (recall that $1/\lambda$ is the average time between rain events). Thus, plants residing in rainy regions, with large $\lambda$ values, have higher discount rate compared to plants that reside in arid regions with low $\lambda$ values. Higher discount rates imply higher myopia, i.e., higher preference of the present over the future, which motivates consumption now over saving for tomorrow. This is perfectly rational: a higher $\lambda$ entails more frequent rain events, hence a higher probability that the next rain event will occur soon, which in turn encourages higher water consumption in the present.

Equation (A.10) represents the expected biomass growth under $t_i < \bar{T}_i$. It holds for all Type 1 management periods and for Type 2 management periods with $t_i < \bar{T}_i$. Equation (A.9) represents the expected biomass growth for Type 2 management periods with $t_i = \bar{T}_i$. For all cases, the optimal transpiration policy given $t_i$ satisfies:

**Proposition 2.** *Given $t_i$, the optimal transpiration policy of Type 1 and Type 2 management periods admits the switching mode*

$$E_i(t) = \begin{cases} \bar{E}(\theta_i(t))\,, & 0 \leq t < \hat{t}_i \\ \underline{E}\,, & \hat{t}_i \leq t \leq t_i \end{cases} \tag{A.11}$$

*for some switching time $\hat{t}_i \in [0, \bar{t}_i]$ chosen such that the SWC process $\theta_i(t)$ that departs from $\theta_i(0)$ and evolves in time according to (A.2) under the switching policy (A.11) reaches $\underline{\theta}$ at $t_i$, i.e.,*

$$\int_0^{\hat{t}_i} [\bar{E}(\theta_i(t)) + D(\theta_i(t))]dt + \int_{\hat{t}_i}^{t_i} [\underline{E} + D(\theta_i(t)]dt = \theta_i(0) - \underline{\theta}\,. \tag{A.12}$$

*Proof.* Given $t_i$, the optimal transpiration policy maximizes

$$\int_0^{t_i} E_i(t)g(m_i(t))e^{-\lambda t}dt$$

subject to (A.2), (A.3), $E_i(t) \in [\underline{E}, \bar{E}(\theta_i(t))]$ and $\theta_i(t_i) = \underline{\theta}$ given the initial states $\theta_i(0)$ and $m_i(0) = m(\tau_{i-1})$. The Hamiltonian corresponding to this problem is

$$\mathcal{H}(t) = e^{-\lambda t}\left\{E_i(t)g(m_i(t)) + \gamma(t)E_i(t)g(m_i(t)) - \eta(t)(E_i(t) + D(\theta_i(t)))\right\}, \tag{A.13}$$

where $\gamma(t)e^{\lambda t}$ and $\eta(t)e^{\lambda t}$ are the costates of $m_i(t)$ and $\theta_i(t)$, respectively. Necessary conditions for optimum include (see, e.g., Kamien and Schwartz 1991):

$$E_i(t) = \begin{cases} \bar{E}(\theta_i(t)) & \text{if } g(m_i(t))(1+\gamma(t)) - \eta(t) > 0 \\ \underline{E} & \text{if } g(m_i(t))(1+\gamma(t)) - \eta(t) < 0 \\ E_i^s(t) & \text{if } g(m_i(t))(1+\gamma(t)) - \eta(t) = 0 \end{cases}, \tag{A.14}$$

where $E_i^s(t)$ is the singular transpiration rate,

$$\dot{\gamma}(t) - \lambda\gamma(t) = -E_i(t)g'(m_i(t))(1+\gamma(t)), \tag{A.15}$$

$$\dot{\eta}(t) - \lambda\eta(t) = \eta(t)D'(\theta_i(t)), \tag{A.16}$$

and the boundary condition

$$\theta_i(t_i) = \underline{\theta}. \tag{A.17}$$

The following Lemma is useful:

**Lemma 2.** *Along the optimal policy,* $\ln(g(m_i(t))(1+\gamma(t))) - \ln(\eta(t))$ *decreases in time.*

*Proof.* Suppressing the time argument and the subscript $i$ for brevity,

$$d(g(m)(1+\gamma))/dt = g'(m)\dot{m}_i(1+\gamma) + g(m)\dot{\gamma}.$$

Substituting for $\dot{m}$ and $\dot{\gamma}$ from (A.3) and (A.15) gives (after some algebraic manipulations)

$$d\ln[g(m)(1+\gamma)]/dt = \lambda\gamma/(1+\gamma).$$

From (A.16) we obtain

$$d\ln(\eta)/dt = \lambda + D'(\theta).$$

It follows that

$$d\left[\ln(g(m)(1+\gamma)) - \ln(\eta)\right]/dt = -\frac{\lambda + (1+\gamma)D'(\theta_i)}{1+\gamma} < 0,$$

completing the proof of the Lemma. □

Back to Proposition 2. As $\theta_i(t)$ contributes to biomass growth, $\eta(t) \geq 0$ and (A.16) ensures that $\eta(t) > 0$. Likewise, $g(m_i(t)) > 0$ and $\gamma(t) \geq 0$ imply $g(m_i(t))(1+\gamma(t)) > 0$. Condition (A.14) can thus be rendered as

$$E_i(t) = \begin{cases} \bar{E}(\theta_i(t)) & \text{if } \ln(g(m_i(t))(1+\gamma(t))) - \ln(\eta(t)) > 0 \\ \underline{E} & \text{if } \ln(g(m_i(t))(1+\gamma(t))) - \ln(\eta(t)) < 0 \\ E_i^s(t) & \text{if } \ln(g(m_i(t))(1+\gamma(t))) - \ln(\eta(t)) = 0 \end{cases}.$$

Lemma 2, then, implies that the optimal $E_i(t)$ admits the switching form (A.11). □

The proposition includes also Type 3 management periods as a special case and verifies Property 1.

The SWC process that departs from $\theta_i(0)$ and evolves in time according to (A.2) under the switching policy (A.11) is denoted $\hat{\theta}_i(t)$. Likewise, the corresponding biomass process that departs from $m_i(0)$ and evolves in time according to (A.3) under the switching policy (A.11) is denoted $\hat{m}_i(t)$.

Equation (A.12) induces the relation $t_i(\hat{t}_i) : [0, \bar{t}_i] \mapsto [\bar{t}_i, \underline{t}_i]$, such that $t_i(\hat{t}_i)$ decreases from $\underline{t}_i$ to $\bar{t}_i$ as $\hat{t}_i$ increases from 0 to $\bar{t}_i$. Writing (A.12) as

$$\int_0^{\hat{t}_i} [\bar{E}(\hat{\theta}_i(t)) + D(\hat{\theta}_i(t))]dt + \int_{\hat{t}_i}^{t_i(\hat{t}_i)} [\underline{E} + D(\hat{\theta}_i(t)]dt = \theta_i(0) - \underline{\theta} \tag{A.18}$$

and differentiating with respect to $\hat{t}_i$ gives

$$\bar{E}(\hat{\theta}_i(\hat{t}_i)) + D(\hat{\theta}_i(\hat{t}_i)) - \left(\underline{E} + D(\hat{\theta}_i(\hat{t}_i)\right) + \left(\underline{E} + D(\hat{\theta}_i(t_i(\hat{t}_i))\right) t_i'(\hat{t}_i) = 0,$$

which in turn implies, recalling that $\hat{\theta}_i(t_i(\hat{t}_i)) = \underline{\theta}$,

$$t_i'(\hat{t}_i) = -\frac{\bar{E}(\hat{\theta}_i(\hat{t}_i)) - \underline{E}}{\underline{E} + D(\underline{\theta})}. \tag{A.19}$$

Now, $\bar{E}(\hat{\theta}_i(\hat{t}_i)) - \underline{E} \geq 0$, equality holding when $\hat{t}_i = \bar{t}_i$. This is so because $\hat{t}_i = \bar{t}_i$ implies $t_i(\bar{t}_i) = \bar{t}_i$, so $\hat{\theta}_i(\hat{t}_i) = \hat{\theta}_i(t_i(\hat{t}_i)) = \underline{\theta}$ and (A.4) implies $\bar{E}(\underline{\theta}) = \underline{E}$. When $\hat{t}_i < \bar{t}_i$, the switching time occurs before $\hat{\theta}_i(t)$ reaches the wilting level $\underline{\theta}$, so $\theta(\hat{t}_i) > \underline{\theta}$ and $\bar{E}(\hat{\theta}_i(\hat{t}_i)) > \underline{E}$. We summarize this discussion in:

**Lemma 3.** *Let $t_i(\hat{t}_i) : [0, \bar{t}_i] \mapsto [\bar{t}_i, \underline{t}_i]$ be defined by (A.18). Then, $t_i'(\hat{t}_i)$, defined in (A.19), satisfies*

$$t_i'(\hat{t}_i) \leq 0 \tag{A.20}$$

*equality holding only at $\hat{t}_i = \bar{t}_i$.*

In light of Proposition 2 and (A.10), the expected biomass growth of Type 1 management periods can be expressed as

$$v_i(\hat{t}_i) = \left(1 - e^{-\lambda t_i(\hat{t}_i)}\right) \left\{ \int_0^{\hat{t}_i} \bar{E}(\hat{\theta}_i(t)) g(\hat{m}_i(t)) e^{-\lambda t} dt + \int_{\hat{t}_i}^{t_i(\hat{t}_i)} \underline{E}\, g(\hat{m}_i(t)) e^{-\lambda t} dt \right\}. \tag{A.21}$$

The optimal transpiration policy is the switching policy (A.11) with the switching time that maximizes $v_i(\hat{t}_i)$:

$$\hat{t}_i^* = \underset{\hat{t}_i \in [0, \bar{t}_i]}{\text{argmax}} \{v(\hat{t}_i)\}. \tag{A.22}$$

It turns out that:

**Proposition 3.** *The condition*

$$\lambda > \underline{E}\,\alpha \tag{A.23}$$

*is sufficient for*

$$\hat{t}_i^* = \bar{t}_i. \tag{A.24}$$

*where $\alpha$ is the water use efficiency (WUE) parameter, i.e., $g(m_i) = \alpha m_i$.*

*Proof.* Differentiating $v_i(\hat{t}_i)$ gives

$$v_i'(\hat{t}_i) = \lambda M_i(\hat{t}_i) e^{-\lambda t_i(\hat{t}_i)} t_i'(\hat{t}_i) + \left(1 - e^{-\lambda t_i(\hat{t}_i)}\right) \times \\ \left\{ \left[ \bar{E}(\hat{\theta}_i(\hat{t}_i)) - \underline{E} \right] g(m_i(\hat{t}_i)) e^{-\lambda \hat{t}_i} + \underline{E}\, g(\hat{m}_i(t_i(\hat{t}_i))) e^{-\lambda t_i(\hat{t}_i)} t_i'(\hat{t}_i) \right\}, \tag{A.25}$$

where

$$M_i(\hat{t}_i) = \int_0^{\hat{t}_i} \bar{E}(\hat{\theta}_i(t)) g(\hat{m}_i(t)) e^{-\lambda t} dt + \int_{\hat{t}_i}^{t_i(\hat{t}_i)} \underline{E}\, g(\hat{m}_i(t)) e^{-\lambda t} dt. \tag{A.26}$$

As $t_i(\bar{t}_i) = \bar{t}_i$ and $\hat{\theta}_i(t_i(\bar{t}_i)) = \hat{\theta}_i(\bar{t}_i) = \underline{\theta}$, Lemma 3 and (A.25) imply

$$v_i'(\bar{t}_i) = 0. \tag{A.27}$$

Moreover, (A.23) ensures

$$v_i'(0) > 0. \tag{A.28}$$

To verify (A.28), use (A.26) and $g(m_i) = \alpha m_i$ to obtain

$$M_i(0) = \int_0^{t_i(0)} \underline{E}\, \alpha \hat{m}_i(t) e^{-\lambda t} dt < \underline{E}\, \alpha \hat{m}_i(t_i(0)) \int_0^{t_i(0)} e^{-\lambda t} dt = \underline{E}\, \alpha \hat{m}_i(t_i(0)) \frac{1}{\lambda} \left(1 - e^{-\lambda t_i(0)}\right).$$

Thus, using (A.19),

$$\lambda M_i(0) e^{-\lambda t_i(0)} [-t_i'(0)] < \left[ \bar{E}(\hat{\theta}(0)) - \underline{E} \right] \frac{\underline{E}}{\underline{E} + D(\underline{\theta})} \alpha \hat{m}_i(t_i(0)) e^{-\lambda t_i(0)} \left(1 - e^{-\lambda t_i(0)}\right). \tag{A.29}$$

Using (A.19) again, the term inside the curely brackets on the right-hand side of (A.25), evaluated at $\hat{t}_i = 0$, becomes

$$\left[ \bar{E}(\hat{\theta}_i(0) - \underline{E} \right] \alpha m_i(0) - \frac{\bar{E}(\hat{\theta}(t_i(0))) - \underline{E}}{\underline{E} + D(\underline{\theta})} \underline{E}\, \alpha \hat{m}_i(t_i(0)) e^{-\lambda t_i(0)} = \\ \left[ \bar{E}(\hat{\theta}_i(0) - \underline{E} \right] \alpha m_i(0) \left[ 1 - \frac{\underline{E}}{\underline{E} + D(\underline{\theta})} \frac{\hat{m}_i(t_i(0))}{\hat{m}_i(0)} e^{-\lambda t_i(0)} \right]. \tag{A.30}$$

Evaluating (A.25) at $\hat{t}_i = 0$ and invoking (A.29)-(A.30) gives

$$v_i'(0) > \left(1 - e^{\lambda t_i(0)}\right)\left[\bar{E}(\hat{\theta}_i(0) - \underline{E}\right]\alpha m_i(0)\left[1 - 2\frac{\underline{E}}{\underline{E} + D(\underline{\theta})}\frac{\hat{m}_i(t_i(0))}{\hat{m}_i(0)}e^{-\lambda t_i(0)}\right]. \tag{A.31}$$

Under the switching policy (A.11) and $\hat{t}_i = 0$, (A.3), recalling $g(m_i) = \alpha m_i$, gives

$$\hat{m}_i(t_i(0)) = \hat{m}_i(0)e^{\underline{E}\alpha t_i(0)},$$

so (A.31) can be rendered as

$$v_i'(0) > \left(1 - e^{\lambda t_i(0)}\right)\left[\bar{E}(\hat{\theta}_i(0) - \underline{E}\right]\alpha m_i(0)\left[1 - 2\frac{\underline{E}}{\underline{E} + D(\underline{\theta})}e^{-[\lambda - \underline{E}\alpha]t_i(0)}\right]. \tag{A.32}$$

It follows that Condition (A.23) ensures that $v_i'(0) > 0$.

Similar, though messier (hence skipped), calculations show that condition (A.23) ensures that $v_i''(\hat{t}_i) < 0$ for $\hat{t}_i \in [0, \bar{t}_i)$. Thus, $v_i'(\hat{t}_i)$ declines from $v_i'(0) > 0$ to $v_i'(\bar{t}_i) = 0$ as $\hat{t}_i$ increases from 0 to $\bar{t}_i$, verifying (A.24). □

Proposition 3 can be rephrased as:

**Proposition 4.** *Under (A.23), the optimal transpiration policy of Type 1 management periods is maximal transpiration throughout.*

Proposition 4 verifies Property 4.

**Remark:** It is verified in Appendix B (see Table B13 and Figure B4) that Condition (A.23) holds for the wild-barley cultivars in all locations considered in this study.

It remains to characterize the optimal switching time of Type 2 management periods. To that end, let $\hat{t}_i^{\bar{T}_i}$ be the switching time satisfying

$$t_i(\hat{t}_i^{\bar{T}_i})) = \bar{T}_i, \tag{A.33}$$

i.e., under the switching policy $\hat{t}_i = \hat{t}_i^{\bar{T}_i}$, the plant reaches maturity at the time the SWC process reaches the wilting level $\underline{\theta}$ (which is feasible for Type 2 management periods). Thus, the policy $\hat{t}_i = \hat{t}_i^{\bar{T}_i}$ guarantees that the plant reaches maturity, in which case Lemma 1 ensures that $\hat{t}_i^* \geq \hat{t}_i^{\bar{T}_i}$. If $\hat{t}_i = \hat{t}_i^{\bar{T}_i}$, the desiccation probability vanishes and the biomass growth equals $M_i(\hat{t}_i^{\bar{T}_i})$. If $\hat{t}_i > \hat{t}_i^{\bar{T}_i}$, the desiccation probability is positive and the expected biomass growth equals $v_i(\hat{t}_i)$, specified in (A.21). Regarding the latter case, invoking Proposition 3 implies that, under (A.23), if $\hat{t}_i^* > \hat{t}_i^{\bar{T}_i}$ then $\hat{t}_i^* = \bar{t}_i$. The reason is that, under maximal transpiration, beyond time $\hat{t}_i^{\bar{T}_i}$ the situation becomes the same as under Type 1 management periods (in which reaching maturity cannot be guaranteed without the help of a rain event even under the minimal transpiration rate). In this case, Proposition 3 implies that maximal transpiration throughout is optimal. Thus, under (A.23), if $\hat{t}_i^* > \hat{t}_i^{\bar{T}_i}$, then $\hat{t}_i^* = \bar{t}_i$. We summarize the above discussion in:

**Proposition 5.** *Under (A.23), the optimal switching time of Type 2 management periods is*

$$\hat{t}_i^* = \begin{cases} \bar{t}_i & \text{if } v_i(\bar{t}_i) \geq M_i(\hat{t}_i^{\bar{T}_i}) \\ \hat{t}_i^{\bar{T}_i} & \text{otherwise} \end{cases} . \tag{A.34}$$

The optimal transpiration during Type 2 management periods begins at the maximal rate and (in the absence of a rain event) lasts until either $\hat{t}_i^{\bar{T}_i}$ or $\bar{t}_i$. In the former case, barring a rain event, the plant switches to minimal transpiration at $\hat{t}_i^{\bar{T}_i}$ and reaches maturity at $\bar{T}_i$. In the latter case, maximal transpiration is applied until the next rain event or desiccation, whatever occurs first. The choice between the two alternatives depends on the relation between $v_i(\bar{t}_i)$ and $M_i(\hat{t}_i^{\bar{T}_i})$, as indicated in (A.34). The proposition verifies Property 3.

#### Summary

- During Type 1 management periods, (A.23) ensures (but is far from necessary) that maximal transpiration throughout is optimal.
- During Type 2 management periods, the optimal transpiration policy is either maximal transpiration until $\hat{t}_i^{\bar{T}_i}$ followed by minimal transpiration until maturity or maximal transpiration throughout. The choice between the two alternatives depends on the comparison between the expected biomass growth under both polices, as indicated in (A.34).
- During Type 3 management periods, maximal transpiration until maturity is optimal.

### References


Kamien, M. I. and Schwartz, N. L.: 1991, *Dynamic optimization: the calculus of variations and optimal control in economics and management*, North-Holland.

Shani, U., Tsur, Y. and Zemel, A.: 2004, Optimal dynamic irrigation schemes, *Optimal Control Applications and Methods* **25**(2), 91–106.
