## Appendix B for "On the biological constraints that limit the productivity of rain-fed annual crops"

### B Empirical Analyses

**Contents**

**B.1 Environmental factors** **2**

B.1.1 Estimation of the expected arrival rate of rainfall events, $\lambda$ 5

B.1.2 NDVI analyses 9

**B.2 Physiological traits** **11**

B.2.1. Length of the growing season ($\bar{T}$) 12

B.2.2. Water-use efficiency ($WUE$) 12

B.2.3. Estimation of maximal and minimal transpiration rates ($\bar{E}$ and $\underline{E}$) 14

**B.3 Empirical analysis of the optimization condition $\boldsymbol{\lambda > WUE \times \underline{E}}$** **28**

[1] Department of Environmental Economics and Management and The Center for Agricultural Economics Research; The Hebrew University of Jerusalem.
[2] Institute of Plant Sciences and Genetics in Agriculture; The Hebrew University of Jerusalem.

### B.1 Environmental factors

This section provides detailed information regarding the estimations of environmental factors related to the model elements, measured for the native sites of the five accessions selected from the B1K wild-barley collection: initial SWC $\theta(0)$, drainage $D$ and rainfall intensity and frequency. The frequency of rain events was measured as the inverse of the expected rainfall arrival rate (denoted $\lambda$, see Appendix A). Geographic, soil and climatic characteristics of the five sites are summarized in Table B1, with the data ordered from the rainiest site in the north of Israel (Meron, with more than 800 mm/year on average) to the driest site in the south (Yeruham, with less than 100 mm/year).

The initial SWC of a management period, $\theta(0)$, increases with the soil's water-retention capacity (e.g., SWC at field capacity), the intensity of the rainfall event at the onset of the management period and the SWC at the end of the previous management period. The latter is expected to increase with rainfall frequency because, all other things being equal, the earlier the next rainfall event arrives, the greater the amount of water remaining in the soil at that point. Soil-water retention capacities are relatively low at the drier southern sites, particularly in Yeruham, where the soil has a higher sand content and higher bulk density than the soils at the other sites. In addition, due to soil-crust effects, the infiltration of rainwater into the soil is particularly low in Yeruham (Eldridge et al. 2000). As shown in Table B1, the average rainfall intensity increases with annual rainfall, as does the frequency of rainfall events. That is, the expected time between subsequent rainfall events is shorter at rainier sites (see Subsection B.1.1 for rainfall-frequency estimation procedures).

**Table B1.** Quantitative measures of environmental factors in the native habitats of the five accessions selected from the B1K wild-barley collection.

| **Item** | **Meron** | **Oren** | **Arbel** | **Guvrin** | **Yeruham** |
|---|---|---|---|---|---|
| B1K code[a] | 51 | 30 | 29 | 35 | 02 |
| **Geographic characteristics** | | | | | |
| X UTM[a] | 726353 | 685269 | 734720 | 680250 | 678260 |
| Y UTM[a] | 3651145 | 3621449 | 3634231 | 3497074 | 3423787 |
| Elevation above sea level (m)[a] | 969 | 98 | 88 | 303 | 535 |
| Slope (degrees)[a] | 5.1 | 13.7 | 19 | 5.2 | 1.6 |
| Aspect (degrees)[a] | 22 | 181 | 48 | 26 | 31 |
| **Soil-structure measures** | | | | | |
| Soil classification[b] | Grumusol / Terra Rossa / Rendzinas | Grumusol / Terra Rossa / Rendzinas | Grumusol / Basaltic proto-grumusol | Dark brown | Lithosols / Loess / Sandy loess |
| Soil bulk density (g $mL^{-1}$)[a] | 1.11 | 1.02 | 1 | 0.97 | 1.41 |
| Soil water content, hygroscopic (%)[a] | 8.17 | 10.38 | 10.82 | 2.91 | 0.54 |
| Soil water content at 1/3 atm (%)[c] | 40.26 | 49.47 | 51.3 | 18.32 | 8.44 |
| **Climatic measures, October–April** | | | | | |
| Daily average temperature (°C)[d] | 12 [D] | 17.2 [AB] | 17.5 [A] | 17.1 [B] | 14.7 [C] |
| Daily average RH (%)[d] | 64.7 [B] | 66.2 [A] | 61.4 [D] | 66.8 [A] | 63.4 [C] |
| Daily average VPD (kPa)[d] | 0.66 [C] | 0.95 [B] | 1.07 [A] | 1.04 [A] | 0.97 [B] |
| Daily average global radiation (W $m^{-2}$)[d] | 348 [C] | 259 [D] | 343 [C] | 383 [B] | 400 [A] |
| Average annual precipitation (mm $year^{-1}$)[d] | 826 [A] | 643 [B] | 449 [C] | 396 [C] | 90 [D] |

| | | | | | |
|---|---|---|---|---|---|
| Average rainfall intensity (mm day$^{-1}$)[d] | 13.6 [A] | 11.9 [B] | 8.6 [C] | 8.3 [C] | 2.4 [D] |
| Expected time between rainfall events (days)[d] | 5.0 [D] | 5.4 [D] | 6.9 [C] | 8.4 [B] | 24.6 [A] |
| **Natural-vegetation indicators** | | | | | |
| Soil organic matter (%)[a] | 8.34 | 9.43 | 7.25 | 5.25 | 0.18 |
| Average NDVI[e] | 0.32 [B] | 0.40 [A] | 0.40 [AB] | 0.43 [AB] | 0.02 [C] |

a. Source: Hübner et al. (2009).

b. Based on Ravikovitch (1992).

c. Computed from hygroscopic water content based on Banin and Amiel (1970).

d. Data are from the meteorological stations closest to each of the five sites (Israel Meteorological Service, (https://ims.gov.il/en). Temperature, relative humidity and VPD data are based on daily averages from 2004 to 2015. Daily global radiation data are 10-min records from October 2016 to April 2017. Daily precipitation data span periods that are specific to each meteorological station. The expected time between rainfall events (i.e., $\frac{1}{\lambda}$, where $\lambda$ is the expected rainfall arrival rate) was obtained from the estimated probability-density functions of rainfall arrival times (see Subsection B.1.1).

e. Data source: Hamaarag (https://www.hamaarag.org.il/en ); averaged NDVI levels were computed for the period December 2013 to April 2014 (see Subsection B.1.2).

One of the factors affecting the drainage rate during management periods ($D$) is soil-water evaporation, which depends on VPD and soil structure. Meron, the site at the highest elevation above sea level, has the lowest daily VPD due to the comparatively low temperatures at that site. The soil bulk density is greatest in Yeruham, implying relatively fast drying out of soil pores at that site (Shokri and Or, 2011).

Soil water is also depleted through its uptake by adjacent plants, which is correlated with vegetation level. An indicator of vegetation is the soil organic-matter content, which is extremely low in Yeruham. Another indication of vegetation is the normalized difference vegetation index (NDVI); the average NDVI levels throughout the barley-growing season are presented in Table B1 (see Section B.1.2). The NDVI data confirm the relatively small amount of vegetation in Yeruham and indicate that barley plants in Meron face less competition for water during the growing season (December to April) than those in Oren, Arbel and Guvrin.

To summarize, both the soil and the climate parameters of the five sites point to a clear north-to-south gradient, in which Meron in the north is a relatively wet and cold site, Yeruham in the south is a desert area, and the habitats of Oren, Arbel and Guvrin are characterized by intermediate environmental conditions.

#### B.1.1 Estimation of the expected arrival rate of rainfall events, $\lambda$

We assume that rainfall events occur at discrete times following a series of Poisson events (see Appendix A). Accordingly, we used the computed frequencies of time intervals between rainfall events to estimate the following probability density function (PDF) of the inter-rainfall time interval $t$

$$PDF(t) = \lambda e^{-\lambda t} \tag{B1}$$

in which $\lambda$ is the rainfall mean arrival rate (number of rain-events per day), implying that $\frac{1}{\lambda}$ (days) is the expected time between subsequent rainfall events.

We obtained daily rainfall data from the Israeli Meteorological Service (https://ims.data.gov.il/). We examined data from the meteorological stations that were closest to each of the five sites represented in the B1K collection. Information on the stations corresponding to each site is presented in Table B2.

**Table B2.** Representative meteorological stations for each of the five B1K sites.

| B1K | **Meron** | **Oren** | **Arbel** | **Guvrin** | **Yeruham** |
|---|---|---|---|---|---|
| Nearest meteorological station | Mount Meron | Bet Oren | Kfar Hitim | Bet Guvrin | Sdeh Boker |
| X UTM | 723262 | 688070 | 734473 | 679565 | 671606 |
| Y UTM | 3655044 | 3623136 | 3631780 | 3499096 | 3416574 |
| Elevation above sea level (m) | 930 | 370 | 40 | 270 | 475 |
| Data period | 1968 – 2007 | 1940 – 2014 | 1945 – 2014 | 1935 – 2014 | 1951 – 2014 |

We consider rainy days as those with at least 5 mm of rainfall. A rain event starts on a rainy day and terminates on the day that is followed by a dry day. We counted the number of days between the end and the beginning of every two consecutive rain events and computed the frequency of each time interval between consecutive rainfall events during the period of data available for each meteorological station.

We estimated the PDF in Equation (B1) for each site by employing a non-linear regression. The results of that estimation are presented in Table B3.

**Table B3.** Estimated $\lambda$ (average number of rain events per day) for the five B1K sites.

| Site | Coef. | Std. Err. | *t* | *P* > *t* | [95% Conf. Interval] | |
|---|---|---|---|---|---|---|
| Meron | 0.198989 | 0.010852 | 18.34 | 0 | 0.177335 | 0.2206428 |
| Oren | 0.185841 | 0.009599 | 19.36 | 0 | 0.166687 | 0.2049944 |
| Arbel | 0.145324 | 0.007432 | 19.55 | 0 | 0.130493 | 0.1601545 |
| Guvrin | 0.118679 | 0.004537 | 26.16 | 0 | 0.109625 | 0.1277322 |
| Yeruham | 0.040652 | 0.003988 | 10.19 | 0 | 0.032694 | 0.048609 |

Goodness-of-fit curves of the estimated PDFs are presented in Fig. B1.

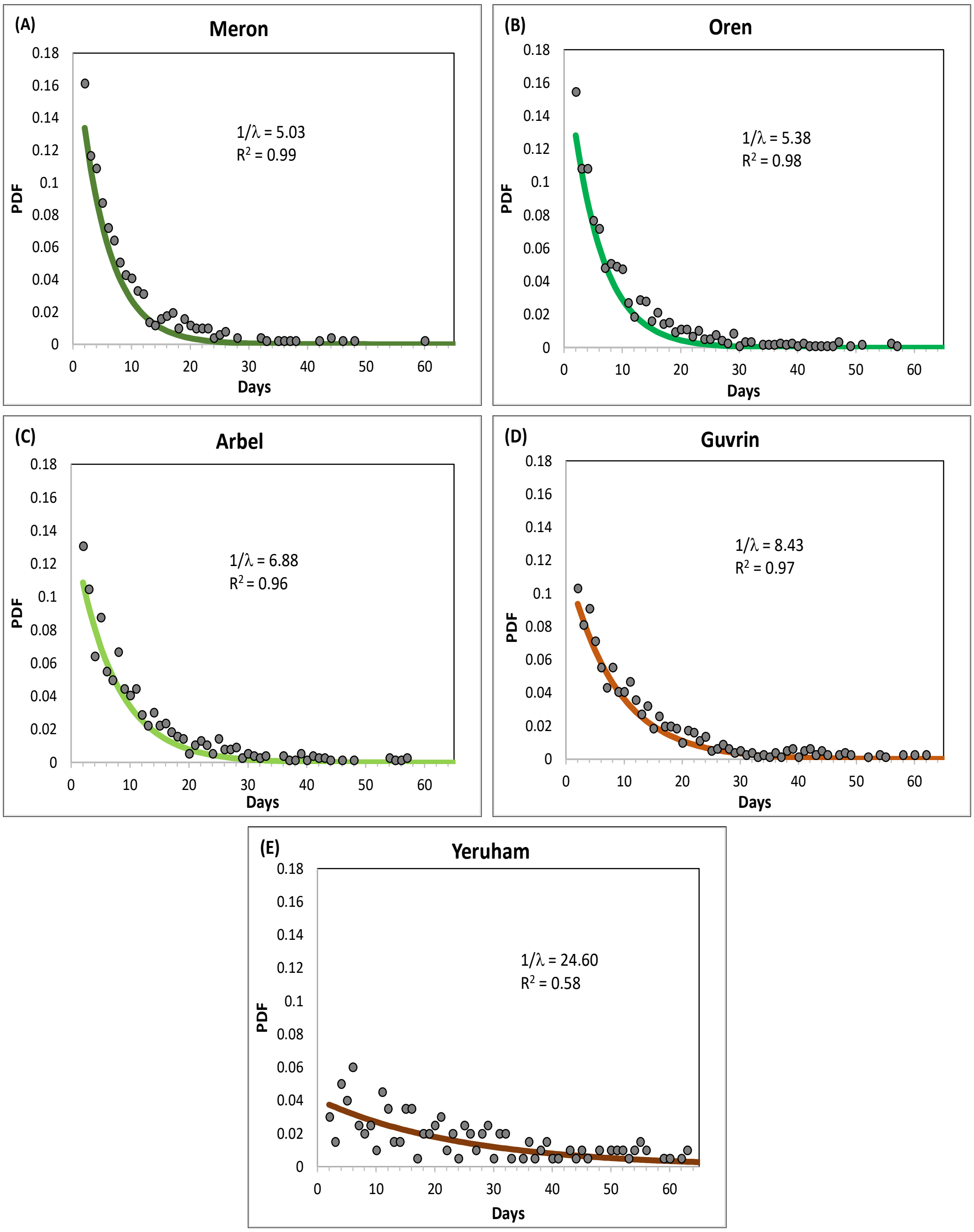

**Fig. B1. Goodness-of-fit curves of the estimated PDFs of inter-rainfall time intervals (Equation B1) for the five B1K sites.** Data source: Israeli Meteorological Service (https://ims.data.gov.il/).

B.1.2 NDVI analyses

NDVI data were provided by Hamaarag, Assessing the State of Israel Nature (http://www.hamaarag.org.il/en) for the five sites, for October 2013 to June 2014, according to the site locations (Table B1). Precipitation data for the same period were taken from the Israeli Meteorological Service (https://ims.data.gov.il/). The levels of variation in NDVI at the five sites are presented in Fig. B2A and the respective cumulative rainfall figures for this period are presented in Fig. B2B.

The level of vegetation in Yeruham was quite low throughout the entire period (Fig. B2A). At the other four sites, vegetation levels increased following a storm during December 2013 (Fig. B2B) and then diminished toward the summer. Yet, the NDVI trajectory in Meron exhibited a relatively slower rate of increase, which peaked in April—about one month later than the maximum NDVI at Oren, Arbel and Guvrin. Thus, if wild barley plants at these four sites germinated after the storm in December 2013, it is likely that those in Meron faced less competition for water during their growing period. This is reflected by the averaged NDVI levels for the period December 2013 to April 2014 (Table B1), which is the typical barley growing season.

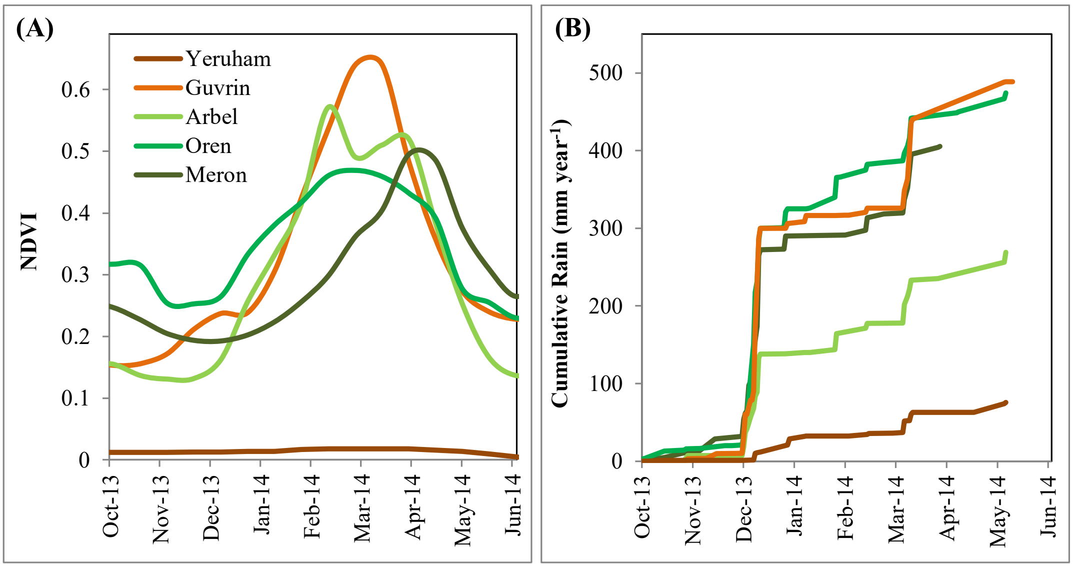

**Fig. B2. (A) NDVI paths and (B) cumulative rainfall from October 2013 to June 2014 at the five B1K sites.** Data sources: Hamaarag, Assessing the State of Israel Nature (http://www.hamaarag.org.il/en); Israeli Meteorological Service (https://ims.data.gov.il/).

### B.2 Physiological traits

This section presents the data and procedures used to derive quantitative measures of the four physiological traits of interest: $WUE$, $\bar{E}$, $\underline{E}$ and $\bar{T}$. The estimated values are reported in Table B4.

**Table B4.** Quantitative traits of the accessions sampled from the five sites selected from the B1K wild-barley collection.

| Item | Meron | Oren | Arbel | Guvrin | Yeruham |
|---|---|---|---|---|---|
| $\hat{t}$ - Mode-switching time (days after planting)[a] | 7.81 $^{A}$ | 7.76 $^{A}$ | 7.85 $^{A}$ | 8.09 $^{A}$ | 7.85 $^{A}$ |
| $\theta_{\hat{t}}$ - Relative SWC at mode switch[a] | 0.15 $^{A}$ | 0.18 $^{A}$ | 0.20 $^{A}$ | 0.22 $^{A}$ | 0.17 $^{A}$ |
| $WUE$ - Water-use efficiency (g $mL^{-1}$)[a] | 0.053 $^{B}$ | 0.056 $^{AB}$ | 0.048 $^{B}$ | 0.062 $^{A}$ | 0.047 $^{B}$ |
| $\bar{E}$ – Maximal transpiration rate = average $E$ in *productive mode*, unstressed plants (mmol $sec^{-1}$ $m^{-2}$)[a] | 3.21 $^{B}$ | 2.45 $^{E}$ | 3.04 $^{C}$ | 2.54 $^{D}$ | 3.24 $^{A}$ |
| $\bar{E}$ – Maximal transpiration rate = average $E$ in *productive mode*, stressed plants (mmol $sec^{-1}$ $m^{-2}$)[a] | 3.55 $^{A}$ | 2.32 $^{D}$ | 2.99 $^{C}$ | 2.20 $^{E}$ | 3.13 $^{B}$ |
| $\underline{E}$ – Minimal transpiration rate = average $E$ in *survival mode*, stressed plants (mmol $sec^{-1}$ $m^{-2}$)[a] | 0.68 $^{A}$ | 0.30 $^{D}$ | 0.48 $^{C}$ | 0.28 $^{E}$ | 0.55 $^{B}$ |
| $\bar{T}$ - length of growing season = flowering time (days from planting)[b] | 66.8 $^{A}$ | 63.6 $^{AB}$ | 61.4 $^{BC}$ | 62.4 $^{AB}$ | 57.8 $^{C}$ |

| | | | | | |
|---|---|---|---|---|---|
| Height at harvest (cm)[b] | 114 AB | 124 A | 121 A | 123 A | 106 C |
| $\bar{m}$ – biomass at maturity = vegetative dry weight at harvest (g)[b] | 57.7 AB | 60.4 AB | 65.3 A | 59.9 AB | 39.6 B |
| Grain size ($cm^2$)[b] | 0.26 A | 0.27 A | 0.29 A | 0.22 B | 0.18 C |
| Grain dry weight (g)[b] | 27.6 A | 23.2 A | 25.7 A | 25.1 A | 12.5 B |

a. Data source: Galkin et al. (2018); see Sections B.2.2 and B.2.3.

b. Data source: Hübner et al. (2013).

B.2.1. Length of the growing season ($\bar{T}$)

For the length of the growing season $\bar{T}$, we relied on data from Hübner et al. (2013), who measured various traits of the B1K collection as they appeared during field experiments conducted under non-water-stressed conditions. In Table B4, we present the average number of days from planting until flowering as a measure of $\bar{T}$, revealing a clear downward trend from the rainiest site to the driest one (See Fig. 7A). Hübner et al. (2013) also reported the vegetative dry weight at harvest—a measure of the biomass at maturity, $\bar{m}$.

B.2.2. Water-use efficiency ($WUE$)

We used output from the PlantArray system water-deficit pot experiments reported by Galkin et al. (2018). Those data incorporate output from four experiments conducted on the five accessions under consideration. Experiments 1 and 2 were conducted in two different greenhouses during December 2012 and Experiments 3 and 4 were conducted during January 2013, in the same greenhouses used for Experiments 1 and 2, respectively.

We estimated $WUE$ based on the data from the non-stressed trials only (i.e., the relative SWC equaled 1 throughout the entire experiment). In these experiments, the plants were irrigated each night from 22:00 to 24:00. We referred to the pot weight at three points of time during the day: 04:00 (after soil-water-plant processes following the irrigation had finished), sunrise and sunset. We defined $Q_{nt}^{kv}$ (g) as the total weight of Pot $n$ of a plant from Accession $k$ ($k = (A, G, M, O, Y)$ for Arbel, Guvrin, Meron, Oren and Yeruham, respectively) and Experiment $v$ ($v = 2, 3, 4$), measured at 04:00 on Day $t$, and $q_{nt}^{kv}$ (g) as the change in the pot's weight between sunrise and sunset on Day $t$. We employed ordinary-least-squares regression to estimate the equation:

$$Q_{nt+1}^{kv} - Q_{nt}^{kv} = (\alpha^k + \alpha^v I_n^v) q_{nt}^{kv} + \varepsilon_{nt}^{kv} \qquad \text{(B2)}$$

where $\alpha^k$ is the $WUE$ coefficient with respect to accession $k$; $I_n^v$ is an indication variable for Experiments 1, 2 and 3; $\alpha^v$ represents the experiment effect and $\varepsilon_{nt}^{kv}$ is the error term. In addition, we clustered observations by pots to allow correlations across standard errors of observations from the same pot.

The results of this estimation are presented in Table B5. The $WUE$ values in Meron, Arbel and Yeruham appear to be relatively small.

**Table B5.** $WUE$ estimates.

| Linear | regression | Number of obs. | = | 1800 |
|---|---|---|---|---|
| | | $F$ (8, 59) | = | 18.61 |
| | | Prob. > $F$ | = | 0 |
| | | $R$-squared | = | 0.0669 |
| | | Root MSE | = | 0.0102 |

(Std. Err. adjusted for 60 clusters in Pot_1)

| Coefficient Symbol | Coefficient Value | Robust Std. Err. | $t$-value | $P > t$ | [95% Conf. | Interval] |
|---|---|---|---|---|---|---|
| $\alpha^A$ | 0.047535 | 0.006202 | 7.67 | 0 | 0.035126 | 0.059945 |
| $\alpha^G$ | 0.061529 | 0.006629 | 9.28 | 0 | 0.048264 | 0.074794 |
| $\alpha^M$ | 0.053205 | 0.006587 | 8.08 | 0 | 0.040025 | 0.066384 |
| $\alpha^O$ | 0.056289 | 0.007168 | 7.85 | 0 | 0.041946 | 0.070631 |
| $\alpha^Y$ | 0.047124 | 0.006115 | 7.71 | 0 | 0.034887 | 0.05936 |
| $\alpha^2$ | -0.04534 | 0.006059 | -7.48 | 0 | -0.05746 | -0.03321 |
| $\alpha^3$ | -0.04295 | 0.006311 | -6.81 | 0 | -0.05558 | -0.03032 |
| $\alpha^4$ | -0.05120 | 0.006692 | -7.65 | 0 | -0.06459 | -0.03781 |

B.2.3. Estimation of maximal and minimal transpiration rates ($\bar{E}$ and $\underline{E}$)

To estimate $\bar{E}$ and $\underline{E}$, we used output from the PlantArray system water-deficit pot experiments reported by Galkin et al. (2018). Our dataset included values of $E$, relative SWC and VPD, all of which were measured at a 3-min resolution in trials with water-stressed and unstressed plants. In each of the water-deficit trials, the $E$ path exhibited a biphasic trajectory, as shown in Fig. 2 and interpreted in Fig. 5 as the *switching mode* optimal-transpiration policy of Type 2 management periods [Property 3(ii)]. In these transpiration trajectories, the maximal transpiration rate $\bar{E}$ was employed (i.e., the *productive mode*) during the first phase, followed by a switch to *survival mode*, in which the minimal transpiration rate $\underline{E}$ was employed. In contrast, non-stressed plants employed $\bar{E}$ throughout the entire experiment.

Data from unstressed plants were used to estimate $\bar{E}$, and to illustrate the $WUE$-$\bar{E}$ constraint $(B1)$ presented in Fig. 6A (as $WUE$ can be estimated only under non-stressed conditions). Stressed-plant trials were the source of the data used to estimate the $\bar{E}$ and $\underline{E}$ levels employed by the plants before and after the mode-switch at time $\hat{t}$, respectively; the estimated values are plotted in Fig. 6B, demonstrating the $\underline{E}$-$\bar{E}$ constraint $(B2)$. Separating the periods of the productive and the survival modes requires the identification of $\hat{t}$, which we defined as the day during which the measured $E$ was most rapidly reduced (e.g., in Fig. 2, it is the 36$^{\text{th}}$ day after planting). Accordingly, the productive-mode period started when irrigation had stopped (i.e., the point at which the relative SWC dropped below 1) and terminated a day before the switching day $\hat{t}$. The survival-mode period began one day after $\hat{t}$ and lasted until the plant was irrigated once again. (All of the plants recovered from the stress, indicating that the SWC did not fall below the wilting-SWC $\underline{\theta}$). The data presented in Table B4 indicate that $\theta_{\hat{t}}$ (the average relative SWC at $\hat{t}$) was comparatively low in plants from Meron and Yeruham; yet we did not obtain statistically significant differences in either $\hat{t}$ or $\theta_{\hat{t}}$ across the different accessions.

With the defined switching day $\hat{t}$, our measures of $\bar{E}$ and $\underline{E}$ were the average levels of $E$ during the periods that the plants were in their productive and survival modes, respectively. However, a simple average of the measured $E$ during each transpiration mode reflects the combined effects of VPD, SWC and the plant's physiological factors (e.g., stomatal aperture); whereas only the latter is of interest for comparison across accessions. Therefore, to identify the differences in $\bar{E}$ and $\underline{E}$ that are attributable to the differences in the accessions' physiological traits, we needed to measure the average level of $E$ during each transpiration mode while neutralizing the impact of the differences in VPD and SWC across trials. To that end, we employed a two-stage estimation procedure: First, we formulated $E$ as a quadratic function of both the VPD during the daytime and relative SWC, taking advantage of the detailed 3-min-resolution data to estimate the function's parameters for each wild-barley accession during each transpiration mode. Then, we used the parameters of the function estimated for a particular accession in a specific mode to compute that accession's predicted $E$ under the average values of VPD and SWC calculated across all of the trials during that mode. For each accession, this procedure yields the value of $E$ computed under similar environmental conditions, such that the remaining differences among accessions reflects only differences in their physiological traits.

For the first estimation stage, we specified quadratic functions for the impact of daytime VPD ($vpd_{nt}^{kv}$) and relative SWC ($\theta_{nt}^{kv}$) of a plant in Pot $n$ from Accession $k$ in Experiment $v$ during daytime of Day $t$ (measured in 3-min intervals) on the maximal and minimal levels of $E$ (i.e., using the observations made during the productive mode and survival mode), respectively:

$$\bar{E}_{nt}^{kv} = vpd_{nt}^{kv}\begin{pmatrix} \alpha_0^k + \alpha_0^v I_n^v + (\alpha_1^k + \alpha_1^v I_n^v)vpd_{nt}^{kv} + \\ (\alpha_2^k + \alpha_2^v I_n^v)\theta_{nt}^{kv} + (\alpha_3^k + \alpha_3^v dI_n^v)(\theta_{nt}^{kv})^2 \end{pmatrix} + \omega_{nt}^{kv} \quad \text{(B3)}$$

$$\underline{E}_{nt}^{kv} = vpd_{nt}^{kv}\begin{pmatrix} \beta_0^k + \beta_0^v d_n^v + (\beta_1^k + \beta_1^v I_n^v)vpd_{nt}^{kv} + \\ (\beta_2^k + \beta_2^v I_n^v)\theta_{nt}^{kv} + (\beta_3^k + \beta_3^v I_n^v)(\theta_{nt}^{kv})^2 \end{pmatrix} + \psi_{nt}^{kv} \quad \text{(B4)}$$

where $\alpha_0^k$ to $\alpha_3^k$, $\alpha_0^v$ to $\alpha_3^v$, $\beta_0^k$ to $\beta_3^k$ and $\beta_0^v$ to $\beta_3^v$ are coefficients and $\omega_{nt}^{kv}$ and $\psi_{nt}^{kv}$ are error terms. For the unstressed-plant experiments (in which $\theta_{nt}^{kv} = 1$ for all values of $t$), we specified the function:

$$\bar{E}_{nt}^{kv} = vpd_{nt}^{kv}\left(\gamma_0^k + \gamma_0^v I_n^v + (\gamma_1^k + \gamma_1^v I_n^v)vpd_{nt}^{kv}\right) + \xi_{nt}^{kv} \quad \text{(B5)}$$

where $\gamma_0^k$, $\gamma_0^v$, $\gamma_1^k$ and $\gamma_1^v$ are parameters and $\xi_{nt}^{kv}$ is the error term. We estimated the functions in Equations (B3), (B4) and (B5) using random effect maximum likelihood (Stata procedure xtreg, mle), while suppressing the constant and clustering observations by pots. The estimation results are reported in Tables B6, B7 and B8 for Equations (B3), (B4) and (B5), respectively. For the second estimation stage, we used the estimated quadratic functions [i.e., Equations (B3), (B4) and (B5)] to predict the values of $\bar{E}$ and $\underline{E}$ under the SWC and VPD averaged across all trials for the respective modes, with standard deviations computed using the bootstrap procedure.

In Table B9, we report the average relative SWC and VPD values used to predict $\bar{E}$ and $\underline{E}$, and compare the average measured values of $\bar{E}$ and $\underline{E}$ and their predicted average values at the average SWC and VPD levels, with SWC and VPD averaged across all trials. The largest difference between the measured and predicted levels is seen with respect to $\underline{E}$ and that difference can be attributed mainly to the differences between the SWC levels that correspond to the survival mode of each accession. For example, the plants from Meron and Yeruham transpired more quickly when they were in productive mode (according to the predictions under similar SWC and VPD conditions), and therefore reached a lower SWC when they switched to the survival mode; consequently, their measured $\underline{E}$ levels (i.e., during the survival mode) were lower than the predicted $\underline{E}$.

Figure B3 is analogous to Fig. 6B, in which $\underline{E}$ is plotted against $\bar{E}$. In Fig. B3, we compare the averaged measured values of $\underline{E}$ versus $\bar{E}$ and their predicted values based on the procedure described above (i.e., under the VPD and SWC averaged across all accessions). Due to the differences in VPD and SWC across the accessions when they were in their productive and survival modes, the correlation (induced by the $\underline{E}$-$\bar{E}$ constraint $B2$) between the measured values of $\underline{E}$ and $\bar{E}$ is much lower than the correlation between their predicted values at the average VPD and SWC levels. Hence, the importance of correcting for differences in VPD and SWC in the application of our proposed breeding procedure (recall Fig. 8).

In Tables B10, B11 and B12, we report the results of the bootstrap analyses conducted for the computation of the predicted values of $\bar{E}$ and $\underline{E}$ for an average Experiment $v$, using the estimated coefficients (Tables B6, B7 and B8, respectively) of Equations (B3), (B4) and (B5), respectively, and the average VPD and SWC (averaged across all accessions; Table B9).

**Table B6.** Estimates of average $\bar{E}$ based on Equation (B3).

| | | | |
|---|---|---|---|
| Random-effects ML regression[a] | Number of obs. | = | 110,330 |
| Group variable: experiment_pot | Number of groups | = | 94 |
| Random effects u_i ~ Gaussian | Obs. per group: | | |
| | Min. | = | 828 |
| | Avg. | = | 1,173.70 |
| | Max. | = | 1,542 |
| | Wald chi2(32) | = | 375544.5 |
| Log likelihood = -169112.6 | Prob > chi2 | = | 0 |

| Coefficient Symbol | Coefficient Value | Std. Err. | *z* | *P* > *z* | [95% Conf. | Interval] |
|---|---|---|---|---|---|---|
| $\alpha_0^A$ | -0.5445 | 0.057472 | -9.47 | 0 | -0.65715 | -0.43186 |
| $\alpha_0^G$ | -0.5038 | 0.064075 | -7.86 | 0 | -0.62938 | -0.37821 |
| $\alpha_0^M$ | -0.60773 | 0.058696 | -10.35 | 0 | -0.72277 | -0.49269 |
| $\alpha_0^O$ | -0.5067 | 0.062099 | -8.16 | 0 | -0.62841 | -0.38499 |
| $\alpha_0^Y$ | -0.69391 | 0.060046 | -11.56 | 0 | -0.8116 | -0.57622 |
| $\alpha_1^A$ | 0.261435 | 0.01343 | 19.47 | 0 | 0.235112 | 0.287758 |
| $\alpha_1^G$ | 0.420458 | 0.013641 | 30.82 | 0 | 0.393722 | 0.447194 |
| $\alpha_1^M$ | 0.397459 | 0.013512 | 29.42 | 0 | 0.370977 | 0.423941 |
| $\alpha_1^O$ | 0.323194 | 0.013735 | 23.53 | 0 | 0.296274 | 0.350114 |
| $\alpha_1^Y$ | 0.383944 | 0.013435 | 28.58 | 0 | 0.357611 | 0.410276 |
| $\alpha_2^A$ | 0.051025 | 0.001408 | 36.23 | 0 | 0.048264 | 0.053785 |

| | | | | | | |
|---|---|---|---|---|---|---|
| $\alpha_2^G$ | 0.016026 | 0.001543 | 10.39 | 0 | 0.013002 | 0.01905 |
| $\alpha_2^M$ | 0.049027 | 0.001424 | 34.44 | 0 | 0.046237 | 0.051817 |
| $\alpha_2^O$ | 0.034279 | 0.001535 | 22.33 | 0 | 0.031271 | 0.037288 |
| $\alpha_2^Y$ | 0.047631 | 0.001415 | 33.65 | 0 | 0.044857 | 0.050405 |
| $\alpha_3^A$ | -0.00024 | 1.15E-05 | -21.35 | 0 | -0.00027 | -0.00022 |
| $\alpha_3^G$ | 4.68E-05 | 1.22E-05 | 3.85 | 0 | 0.000023 | 7.06E-05 |
| $\alpha_3^M$ | -0.00018 | 1.15E-05 | -15.8 | 0 | -0.0002 | -0.00016 |
| $\alpha_3^O$ | -0.00015 | 1.23E-05 | -11.78 | 0 | -0.00017 | -0.00012 |
| $\alpha_3^Y$ | -0.0002 | 1.14E-05 | -17.51 | 0 | -0.00022 | -0.00018 |
| $\alpha_0^2$ | 0.164775 | 0.063761 | 2.58 | 0.01 | 0.039806 | 0.289745 |
| $\alpha_0^3$ | 0.90289 | 0.065089 | 13.87 | 0 | 0.775318 | 1.030461 |
| $\alpha_0^4$ | -0.26721 | 0.071825 | -3.72 | 0 | -0.40799 | -0.12644 |
| $\alpha_1^2$ | -0.24118 | 0.022417 | -10.76 | 0 | -0.28511 | -0.19724 |
| $\alpha_1^3$ | -0.41284 | 0.013037 | -31.67 | 0 | -0.4384 | -0.38729 |
| $\alpha_1^4$ | -0.28632 | 0.014031 | -20.41 | 0 | -0.31382 | -0.25882 |
| $\alpha_2^2$ | 0.055885 | 0.001424 | 39.24 | 0 | 0.053093 | 0.058676 |
| $\alpha_2^3$ | 0.004254 | 0.001526 | 2.79 | 0.005 | 0.001264 | 0.007245 |
| $\alpha_2^4$ | 0.022459 | 0.001683 | 13.35 | 0 | 0.019161 | 0.025757 |
| $\alpha_3^2$ | -0.00043 | 1.21E-05 | -35.66 | 0 | -0.00045 | -0.00041 |
| $\alpha_3^3$ | -0.00012 | 1.19E-05 | -9.9 | 0 | -0.00014 | -9.4E-05 |
| $\alpha_3^4$ | -0.00015 | 1.31E-05 | -11.55 | 0 | -0.00018 | -0.00013 |
| /sigma_u | 0.699892 | 0.051408 | | | 0.606051 | 0.808264 |
| /sigma_e | 1.117664 | 0.00238 | | | 1.113008 | 1.122339 |
| rho | 0.281681 | 0.029737 | | | 0.226479 | 0.342659 |

a. LR test of sigma_u = 0: chibar2(01) = 2.5e+04      Prob >= chibar2 = 0.000.

**Table B7.** Estimates of average $\underline{E}$ based on Equation (B4).

| | | | |
|---|---|---|---|
| Random-effects ML regression[a] | Number of obs. | = | 160654 |
| Group variable: experiment_pot | Number of groups | = | 92 |
| Random effects u_i ~ Gaussian | Obs. per group: | | |
| | Min. | = | 218 |
| | Avg. | = | 1746.2 |
| | Max. | = | 2889 |
| | Wald chi2(32) | = | 13106.56 |
| Log likelihood = -302516.84 | Prob > chi2 | = | 0 |

| Coefficient Symbol | Coefficient Value | Std. Err. | *z* | *P* > *z* | [95% Conf. Interval] | |
|---|---|---|---|---|---|---|
| $\beta_0^A$ | -0.11699 | 0.09911 | -1.18 | 0.238 | -0.31124 | 0.077263 |
| $\beta_0^G$ | -0.60851 | 0.102141 | -5.96 | 0 | -0.80871 | -0.40832 |
| $\beta_0^M$ | -0.31209 | 0.097679 | -3.2 | 0.001 | -0.50354 | -0.12065 |
| $\beta_0^O$ | -0.5779 | 0.101045 | -5.72 | 0 | -0.77595 | -0.37986 |
| $\beta_0^Y$ | -0.42576 | 0.101551 | -4.19 | 0 | -0.6248 | -0.22672 |
| $\beta_1^A$ | -0.08288 | 0.038716 | -2.14 | 0.032 | -0.15876 | -0.007 |
| $\beta_1^G$ | 0.098963 | 0.039846 | 2.48 | 0.013 | 0.020867 | 0.177059 |
| $\beta_1^M$ | 0.01116 | 0.037954 | 0.29 | 0.769 | -0.06323 | 0.085548 |
| $\beta_1^O$ | 0.073778 | 0.039164 | 1.88 | 0.06 | -0.00298 | 0.150538 |
| $\beta_1^Y$ | 0.012638 | 0.038905 | 0.32 | 0.745 | -0.06362 | 0.088891 |
| $\beta_2^A$ | 0.076594 | 0.005434 | 14.1 | 0 | 0.065944 | 0.087245 |
| $\beta_2^G$ | 0.066607 | 0.004695 | 14.19 | 0 | 0.057406 | 0.075809 |
| $\beta_2^M$ | 0.074567 | 0.006438 | 11.58 | 0 | 0.061948 | 0.087186 |
| $\beta_2^O$ | 0.076879 | 0.005569 | 13.8 | 0 | 0.065963 | 0.087794 |
| $\beta_2^Y$ | 0.088147 | 0.006274 | 14.05 | 0 | 0.075851 | 0.100443 |

| | | | | | | |
|---|---|---|---|---|---|---|
| $\beta_3^A$ | -0.00132 | 0.000133 | -9.93 | 0 | -0.00158 | -0.00106 |
| $\beta_3^G$ | -0.0007 | 6.99E-05 | -10 | 0 | -0.00084 | -0.00056 |
| $\beta_3^M$ | -0.00021 | 0.000218 | -0.98 | 0.327 | -0.00064 | 0.000213 |
| $\beta_3^O$ | -0.00112 | 0.000136 | -8.27 | 0 | -0.00139 | -0.00086 |
| $\beta_3^Y$ | -0.0009 | 0.000179 | -5.04 | 0 | -0.00125 | -0.00055 |
| $\beta_0^2$ | 0.326623 | 0.121855 | 2.68 | 0.007 | 0.087791 | 0.565455 |
| $\beta_0^3$ | 0.494056 | 0.097112 | 5.09 | 0 | 0.30372 | 0.684392 |
| $\beta_0^4$ | 0.123548 | 0.098312 | 1.26 | 0.209 | -0.06914 | 0.316235 |
| $\beta_1^2$ | -0.06954 | 0.052313 | -1.33 | 0.184 | -0.17207 | 0.032995 |
| $\beta_1^3$ | -0.07572 | 0.037657 | -2.01 | 0.044 | -0.14953 | -0.00191 |
| $\beta_1^4$ | -0.07348 | 0.038674 | -1.9 | 0.057 | -0.14928 | 0.002324 |
| $\beta_2^2$ | -0.06274 | 0.013229 | -4.74 | 0 | -0.08867 | -0.03682 |
| $\beta_2^3$ | -0.0535 | 0.005239 | -10.21 | 0 | -0.06377 | -0.04324 |
| $\beta_2^4$ | 0.005311 | 0.005458 | 0.97 | 0.331 | -0.00539 | 0.016008 |
| $\beta_3^2$ | 0.003375 | 0.000831 | 4.06 | 0 | 0.001747 | 0.005004 |
| $\beta_3^3$ | 0.001231 | 0.00013 | 9.48 | 0 | 0.000977 | 0.001486 |
| $\beta_3^4$ | 0.000413 | 0.00013 | 3.17 | 0.002 | 0.000158 | 0.000669 |
| /sigma_u | 0.29605 | 0.02511 | | | 0.250709 | 0.349592 |
| /sigma_e | 1.588779 | 0.002804 | | | 1.583293 | 1.594284 |
| rho | 0.033557 | 0.005503 | | | 0.02411 | 0.045859 |

a. Likelihood-ratio test of sigma_u = 0: chibar2(01) = 1692.85 Prob > =chibar2 = 0.000.

**Table B8.** Estimates of average $\bar{E}$ based on Equation (B5).

| | | | |
|---|---|---|---|
| Random-effects ML regression[a] | Number of obs. | = | 351,656 |
| Group variable: pot_exp | Number of groups | = | 60 |
| Random effects u_i ~ Gaussian | Obs. per group: | | |

| | | | |
|---|---|---|---|
| | Min. | = | 3,797 |
| | Avg. | = | 5,860.90 |
| | Max. | = | 7,697 |
| | Wald chi2(16) | = | 91188.53 |
| Log likelihood = -918028.7 | Prob > chi2 | = | 0 |

| Coefficient Symbol | Coefficient Value | Std. Err. | $z$ | $P > z$ | [95% Conf. Interval] | |
|---|---|---|---|---|---|---|
| $\gamma_0^A$ | 2.995322 | 0.046987 | 63.75 | 0 | 2.903229 | 3.087416 |
| $\gamma_0^G$ | 2.133129 | 0.046942 | 45.44 | 0 | 2.041125 | 2.225133 |
| $\gamma_0^M$ | 2.815571 | 0.047779 | 58.93 | 0 | 2.721926 | 2.909216 |
| $\gamma_0^O$ | 2.173696 | 0.046995 | 46.25 | 0 | 2.081588 | 2.265803 |
| $\gamma_0^Y$ | 2.767333 | 0.047567 | 58.18 | 0 | 2.674104 | 2.860563 |
| $\gamma_1^A$ | -0.31399 | 0.020196 | -15.55 | 0 | -0.35358 | -0.27441 |
| $\gamma_1^G$ | -0.05885 | 0.020191 | -2.91 | 0.004 | -0.09842 | -0.01927 |
| $\gamma_1^M$ | -0.13825 | 0.02049 | -6.75 | 0 | -0.17841 | -0.09809 |
| $\gamma_1^O$ | -0.12218 | 0.020194 | -6.05 | 0 | -0.16176 | -0.0826 |
| $\gamma_1^Y$ | -0.10122 | 0.020461 | -4.95 | 0 | -0.14132 | -0.06111 |
| $\gamma_0^2$ | 0.677354 | 0.053074 | 12.76 | 0 | 0.573331 | 0.781376 |
| $\gamma_0^3$ | -1.57032 | 0.039389 | -39.87 | 0 | -1.64752 | -1.49312 |
| $\gamma_0^4$ | -0.14177 | 0.042062 | -3.37 | 0.001 | -0.22421 | -0.05933 |
| $\gamma_1^2$ | 0.214035 | 0.029065 | 7.36 | 0 | 0.157067 | 0.271002 |
| $\gamma_1^3$ | 0.226194 | 0.01793 | 12.62 | 0 | 0.191052 | 0.261337 |
| $\gamma_1^4$ | -0.11794 | 0.019418 | -6.07 | 0 | -0.156 | -0.07988 |
| /sigma_u | 0.666025 | 0.06165 | | | 0.55552 | 0.798511 |
| /sigma_e | 3.290973 | 0.003925 | | | 3.28329 | 3.298674 |
| rho | 0.039346 | 0.006998 | | | 0.027452 | 0.055131 |

a. LR test of sigma_u = 0: chibar2(01) = 7952.04 Prob > = chibar2 = 0.000.

**Table B9.** Average measures of $\overline{E}$ and $\underline{E}$ versus their predicted values calculated using Equations (B3), (B4) and (B5) at the average VPD and SWC across all accessions in the productive and survival modes, respectively.

| | VPD (kPa) | SWC (%) | Measured (mmol $sec^{-1}$ $m^{-2}$) | Predicted under average VPD and SWC (mmol $sec^{-1}$ $m^{-2}$) |
|---|---|---|---|---|
| **$\overline{E}$, unstressed plants** | | | | |
| Meron | 1.24 | 100 | 3.20 | 3.21 |
| Oren | 1.27 | 100 | 2.43 | 2.45 |
| Arbel | 1.27 | 100 | 2.91 | 3.04 |
| Guvrin | 1.25 | 100 | 2.63 | 2.54 |
| Yeruham | 1.23 | 100 | 3.24 | 3.24 |
| Average | 1.25 | 100 | | |
| **$\overline{E}$, stressed plants** | | | | |
| Meron | 1.51 | 61.37 | 3.84 | 4.08 |
| Oren | 1.49 | 63.33 | 2.90 | 2.85 |
| Arbel | 1.49 | 63.14 | 3.32 | 3.52 |
| Guvrin | 1.50 | 65.98 | 2.79 | 2.73 |
| Yeruham | 1.51 | 62.97 | 3.65 | 3.65 |
| Average | 1.50 | 63.30 | | |
| **$\underline{E}$, stressed plants** | | | | |
| Meron | 1.12 | 7.71 | 0.42 | 0.68 |

| | | | | |
|---|---|---|---|---|
| Oren | 1.12 | 10.21 | 0.41 | 0.30 |
| Arbel | 1.14 | 15.72 | 0.60 | 0.48 |
| Guvrin | 1.12 | 11.37 | 0.36 | 0.28 |
| Yeruham | 1.11 | 9.33 | 0.44 | 0.55 |
| Average | 1.12 | 10.86 | | |

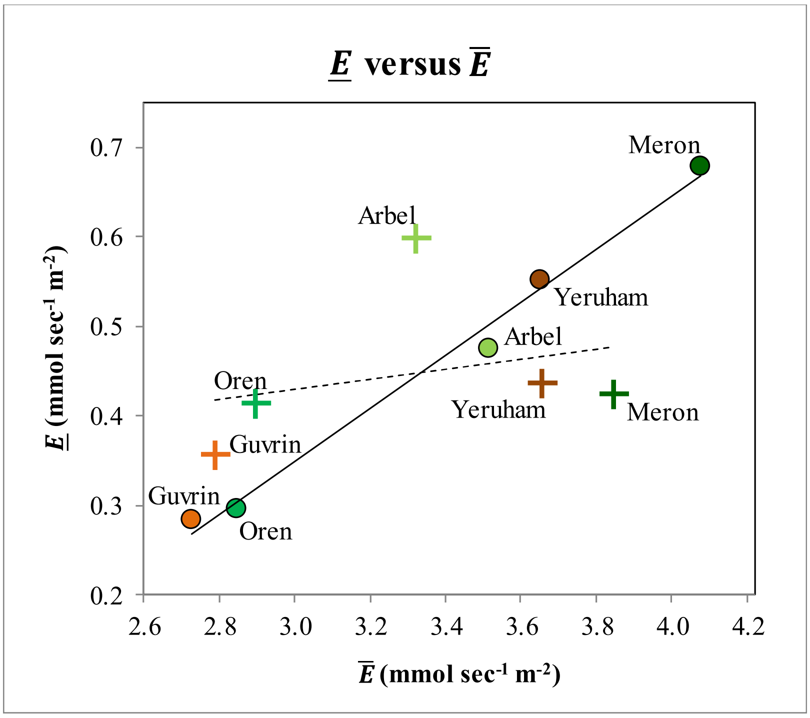

**Fig. B3. $\underline{E}$ plotted against $\bar{E}$ (both estimated for water-stressed plants).** The plus (+) signs represent the measured values and the round dots (o) are the predicted values under the VPD and SWC averaged across all accessions (Table B9). The correlation between the $\underline{E}$ and $\bar{E}$ predicted under similar VPD and SWC (solid linear trend line) is larger than that of the measured $\underline{E}$ and $\bar{E}$ values (dashed linear trend line).

**Table B10.** Results of bootstrap analyses conducted to predict $\bar{E}$ (mmol sec$^{-1}$ m$^{-2}$) for an average Experiment $v$, using the estimated coefficients (Table B6) of Equation (B3) and the VPD and SWC averaged across all accessions (Table B9).

| Site | Coef. | Std. Err. | $z$ | $P > z$ | [95% Conf. Interval] | |
|---|---|---|---|---|---|---|
| Meron | 4.076355 | 0.020583 | 181.16 | 0 | 3.688509 | 3.769194 |
| Oren | 2.848999 | 0.018326 | 136.5 | 0 | 2.465621 | 2.537457 |
| Arbel | 3.516129 | 0.021695 | 146.06 | 0 | 3.126177 | 3.211219 |
| Guvrin | 2.728448 | 0.021016 | 113.28 | 0 | 2.339592 | 2.421973 |
| Yeruham | 3.651162 | 0.01919 | 172.15 | 0 | 3.265849 | 3.34107 |

**Table B11.** Results of bootstrap analyses conducted for predicting $\underline{E}$ (mmol sec$^{-1}$ m$^{-2}$) for an average Experiment $v$, using the estimated coefficients (Table B7) of Equation (B4) and the VPD and SWC averaged across all accessions (Table B9).

| Site | Coef. | Std. Err. | $z$ | $P > z$ | [95% Conf. Interval] | |
|---|---|---|---|---|---|---|
| Meron | 0.678256 | 0.0148 | 36.82 | 0 | 0.516077 | 0.574105 |
| Oren | 0.295764 | 0.0120 | 13.56 | 0 | 0.139094 | 0.186086 |
| Arbel | 0.475457 | 0.0136 | 25.13 | 0 | 0.315596 | 0.368996 |
| Guvrin | 0.284715 | 0.0126 | 12.06 | 0 | 0.126914 | 0.176159 |
| Yeruham | 0.551186 | 0.0146 | 28.72 | 0 | 0.389495 | 0.446552 |

**Table B12.** Results of bootstrap analyses conducted for predicting $\bar{E}$ (mmol sec$^{-1}$ m$^{-2}$) for an average Experiment $v$, using the estimated coefficients (Table B8) of Equation (B5) and the VPD and SWC averaged across all accessions (Table B9).

| Site | Coef. | Std. Err. | $z$ | $P > z$ | [95% Conf. Interval] | |
|---|---|---|---|---|---|---|
| Meron | 3.214325 | 0.016703 | 192.44 | 0 | 3.181587 | 3.247062 |
| Oren | 2.446686 | 0.015486 | 157.99 | 0 | 2.416335 | 2.477038 |
| Arbel | 3.042843 | 0.01752 | 173.68 | 0 | 3.008505 | 3.077182 |
| Guvrin | 2.5388 | 0.017112 | 148.37 | 0 | 2.505262 | 2.572338 |
| Yeruham | 3.237484 | 0.016047 | 201.76 | 0 | 3.206034 | 3.268935 |

**B.3 Empirical analysis of the optimization condition $\lambda > WUE \times \underline{E}$**

According to Proposition 4, the inequality [see Eq. (A.23) in Appendix A]:

$$\lambda > WUE \times \underline{E} \tag{B6}$$

is a sufficient (but not necessary) condition for the optimality of the productive-mode transpiration policy throughout the entire Type 1 management period (during which period, survival until maturity is beyond reach). That is, if the condition is met, the plant should continuously employ the most risky transpiration rate, as the SWC declines most rapidly toward the wilting level $\underline{\theta}$, unless rain arrives before that SWC level is reached.

Let us interpret condition (B6). The units of both sides are time$^{-1}$. Considering the right-hand side, $WUE$ stands for the biomass accumulated per unit of transpired water and $\underline{E}$ is the water transpired per unit of time, divided by the accumulated biomass. Therefore, the term $WUE \times \underline{E}$ is the benefit (in terms of relative biomass per unit of time) obtained if the plant minimizes its desiccation risk. Regarding the left-hand side, let us rewrite it as follows:

$$\lambda \times \frac{biomass\ survived\ if\ rainfall\ arrives}{accumulated\ biomass} \tag{B7}$$

where $\lambda$ (the rainfall mean arrival rate) is multiplied by the term $\frac{biomass\ survived\ if\ rainfall\ arrives}{accumulated\ biomass}$, which is unit-less and equals 1 (because the surviving biomass in a situation in which rain arrives equals the accumulated biomass). Accordingly, (B7) is the expected benefit (in terms of relative biomass per unit of time) if the plant survives due to the arrival of rain. Thus, Condition (B6) implies that, throughout a Type 1 management period, the expected benefit associated with survival through the arrival of rainfall is greater than the benefit associated with minimizing the desiccation risk. Therefore, switching to the minimal transpiration rate is suboptimal, which [given Proposition 2, see Eq. (A11) in Appendix A] implies that transpiring at the highest rate throughout the entire management period is the optimal behavior.

Our objective in this section is to test whether Condition (B6) held true for our five barley accessions. In Table B13, we present the estimated values of the product $WUE \times \underline{E}$ and $\lambda$. These data indicate that the condition is indeed met with respect to the accessions from all sites. In Fig. B4, we present an illustration of the comparison between $WUE \times \underline{E}$ and the rainfall mean arrival rate $\lambda$.

**Table B13.** Estimated values of $\underline{E}$ (mL $g^{-1}$ $day^{-1}$), $WUE$ (g $mL^{-1}$), $WUE \times \underline{E}$ ($day^{-1}$) and $\lambda$ ($day^{-1}$) in relation to Condition (B6).

| **Site** | $\underline{\boldsymbol{E}}$ | $\boldsymbol{WUE}$ | $\boldsymbol{WUE \times \underline{E}}^{b}$ | $\boldsymbol{\lambda}^{c}$ |
|---|---|---|---|---|
| Meron | 0.598<br>(0.013) | 0.053<br>(0.012) | 0.032<br>(0.007) | 0.199<br>(0.011) |
| Oren | 0.283<br>(0.011) | 0.056<br>(0.013) | 0.016<br>(0.004) | 0.186<br>(0.01) |
| Arbel | 0.365<br>(0.01) | 0.048<br>(0.013) | 0.017<br>(0.005) | 0.145<br>(0.007) |
| Guvrin | 0.325<br>(0.014) | 0.062<br>(0.013) | 0.02<br>(0.004) | 0.119<br>(0.005) |

| | | | | |
|---|---|---|---|---|
| Yeruham | 0.557<br>(0.015) | 0.047<br>(0.012) | 0.026<br>(0.007) | 0.041<br>(0.004) |

a. Numbers in brackets are standard errors.

b. Since $\underline{E}$ and $WUE$ are estimated based on different experiments, their covariance is unknown. Therefore, we computed the standard errors of the product $WUE \times \underline{E}$ based on the assumption that $\underline{E}$ and $WUE$ are independent.

c. Taken from Section B.1.1.

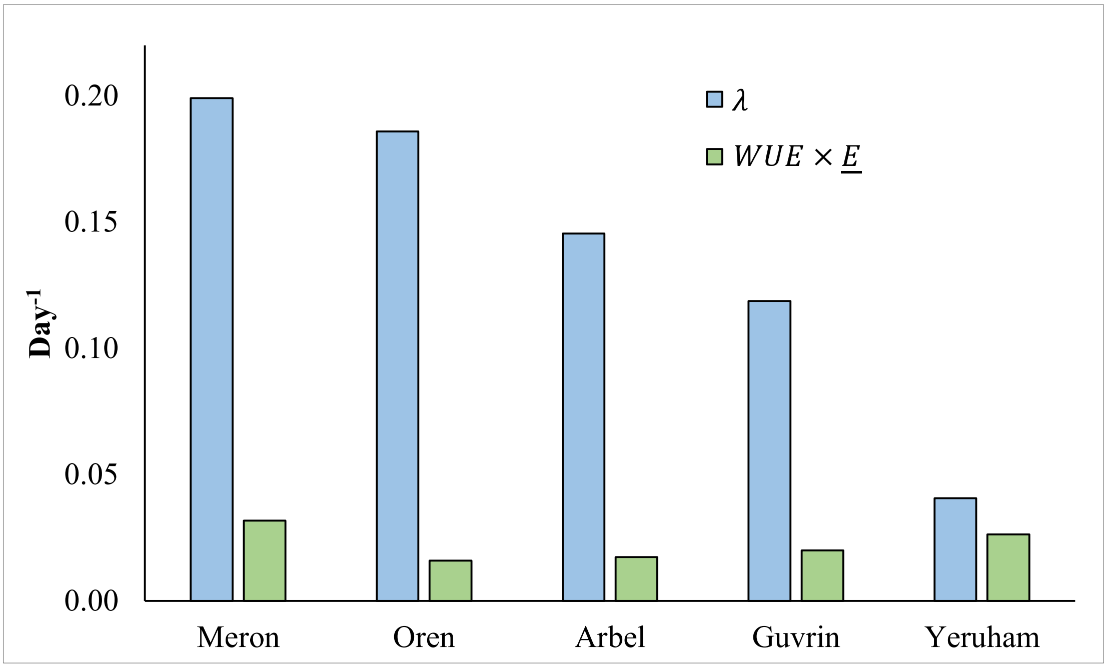

**Fig. B4. Estimated values of $\lambda$ and $WUE \times \underline{E}$ in relation to Condition** (B6).

We mention three points with respect to the empirical test whose results are reported in Table B13 and Fig. B4. First, the estimations of $\underline{E}$ and $WUE$ were based on pot experiments in which all plants were grown under similar soil and atmospheric conditions, which differ from those in the plants' natural habitats. (This is because those experiments aimed to compare the accessions under similar conditions.) While $\underline{E}$ and $WUE$, as defined by the theoretical model, should not depend on environmental conditions, their experimentally measured values may. Second, our estimation of $\lambda$ is based on precipitation data from a few recent decades (Appendix B.1.1) that represent the current climates at the sites of interest. However, from the perspective of evolution, one should consider the climate over millions of years. Climate records indicate that the present climate is nearly 6°C warmer than the average temperature during the last 800,000 years (Lüthi et al. 2008). This implies that the long-term average levels of $\lambda$ at our five sites are likely greater than those reported in Table B13 and Fig. B4, which strengthens our conclusion that Condition (B6) is satisfied. Finally, the lower the value of $\underline{E}$, the longer the plant can survive during a management period and, therefore, Type 1 periods fall earlier in the growing season, when the plant is relatively small. Since experimental analyses of water management by small plants is challenging, so is the verification of Proposition 4 by experimental observation.

**References**

Banin, A., & Amiel, A. (1970). A correlative study of the chemical and physical properties of a group of natural soils of Israel. *Geoderma*, *3*(3), 185-198.

Galkin, E., Dalal, A., Evenko, A., Fridman, E., Kan, I., Wallach, R. & Moshelion, M. (2018). Risk-management strategies and transpiration rates of wild barley in uncertain environments. *Physiologia Plantarum*, *164*(4), 412-428.

Hübner, S., Hoffken, M., Oren, E., Haseneyer, G., Stein, N., Graner, A., Schmid, K. & Fridman, E. (2009). Strong correlation of wild barley (*Hordeum spontaneum*) population structure with temperature and precipitation variation. *Molecular Ecology*, *18*, 1523-1536.

Hübner, S., Bdolach, E., Ein-Gedy, S., Schmid, K. J., Korol, A., & Fridman, E. (2013). Phenotypic landscapes: phenological patterns in wild and cultivated barley. *Journal of Evolutionary Biology*, *26*(1), 163-174.

Lüthi, D., Le Floch, M., Bereiter, B., Blunier, T., Barnola, J. M., Siegenthaler, U., Raynaud, D., Jouzel, J., Fischer, H., Kawamura, K. & Stocker, T. F. (2008). High-resolution carbon dioxide concentration record 650,000–800,000 years before present. *nature*, *453*(7193), 379-382.

Ravikovitch, S. (1992). The soils of Israel: Formation, nature, and properties, Bnei Brak: Hakibbuts Hamehuchad Publishing (Hebrew).

Shokri, N. and Or, D. (2011). What determines drying rates at the onset of diffusion controlled stage-2 evaporation from porous media?. *Water Resources Research*, *47*(9).
